# Contrasting responses of protistan plant parasites and phagotrophs to ecosystems, land management and soil properties

**DOI:** 10.1101/2020.03.22.001768

**Authors:** Anna Maria Fiore-Donno, Tim Richter-Heitmann, Michael Bonkowski

## Abstract

Functional traits are increasingly used in ecology to link the structure of microbial communities to ecosystem processes. We investigated two important protistan lineages, Cercozoa and Endomyxa (Rhizaria) in soil using Illumina sequencing and analysed their diversity and functional traits along with their responses to environmental factors in grassland and forest across Germany. From 600 soil samples, we obtained 2,101 Operational Taxonomy Units representing ~18 million Illumina reads (region V4, 18S rRNA gene). All major taxonomic and functional groups were present, dominated by small bacterivorous flagellates (Glissomonadida). Endomyxan plant parasites were absent from forest. In grassland, they were promoted by more intensive land use management. Grassland and forest strikingly differed in community composition. Relative abundances of bacterivores and eukaryvores were contrastingly influenced by environmental factors, indicating bottom-up regulation by food resources. These patterns provide new insights into the functional organization of soil biota and indications for a more sustainable land-use management.

**Highlights:**

- Protistan plant parasites of worldwide importance (Phytomyxea) are absent from forest
- Protistan plant parasites are enhanced by land use intensification in grassland
- Opposite responses of protistan trophic guilds to environmental conditions in forest
- Drastic differences in protistan community composition between grassland and forest

## 1. Introduction

A major aim in ecology is to understand the drivers affecting the composition and functioning of the soil food web, and how its components contribute to ecosystem functions and services. Because soil biota are incredibly diverse, they are commonly aggregated into feeding groups (guilds or trophic species) to facilitate the understanding on their complex interactions (Scheu, 2002). Protists make up a non-negligible fraction (30-40%) of the soil eukaryotes (Shen et al., 2014; Lanzén et al., 2016; Ferreira de Araujo et al., 2018), and they have long been recognized as a pivotal component of soil food webs (Hunt et al., 1987). Despite of this, they have only recently been included in general models on ecosystem services (de Vries et al., 2014), probably because of their immense diversity. However, uncertainties in consumers’ preferences in soil food webs can strongly influence model predictions (Reyns et al., 2019). A main obstacle in food web models is the traditional simplistic, but erroneous assumption that soil protists act only as bacterivores - instead protists occupy all trophic levels, and include, next to bacterivores, autotrophs, mixotrophs, saprotrophs, eukaryvores, omnivores, as well as parasites of animals and plants and their hyperparasites (Geisen et al., 2016; Bonkowski et al., 2019). This diversity of trophic roles of protists fundamentally affects the efficiency of the energy flow across the soil food web, which does not follow a single-track channel from small to large consumers as traditionally assumed (Potatov et al., 2019). Disentangling the multiple roles of protists will thus increase the trophic resolution of the soil food web and will have a major influence on network properties, such as topology and connectivity (Henriksen et al., 2019). Therefore, it is essential, not only to conduct large-scale molecular environmental sampling studies in soil, but also to attribute ecologically meaningful traits to the identified sequences. This is now facilitated by publicly available databases of protistan functional traits, including trophic guilds (Bjorbækmo et al., 2019; Dumack et al., 2019), and by community efforts to improve the reference databases for taxonomic annotation (del Campo et al., 2018).

Recent large-scale environmental sampling investigations of soil protists in natural and semi-natural environments gave insights into their multiple feeding modes, and how each trophic guild reacts to environmental filters and human-induced disturbances. Nonetheless, it must be noted that although the relative proportion of the main trophic guilds is known for planktonic protists (43 % symbionts, 43 % predators) (Bjorbækmo et al., 2019), such a comprehensive catalogue has not yet been realized for soil. Only inventories limited - in space, by their specific aims and by methodological biases - gave partially overlapping results: a worldwide soil survey indicated that protistan communities were largely composed of “consumers” with a minority of parasites and phototrophs (Oliverio et al., 2020). More precisely, in a single grassland soil tackling specific lineages, “consumers” were composed of bacterivores (67%), while omnivores (feeding on bacteria and other eukaryotes) and eukaryvores together reached a non-negligible 25% (Fiore-Donno et al., 2019). Even less is known about the distribution of protistan plant pathogens in natural habitats: most of the protists that are known to interact with plants belong to the Stramenopiles-Alveolata-Rhizaria super group (Harosa or “SAR”), particularly those belonging to oomycetes (Stramenopiles) and Cercozoa (Rhizaria) (Hassani et al., 2018). For example, Schulz et al. (2019), investigating a gradient of land use intensification from tropical rain forest to plantations, reported significant shifts in the composition of protistan phagotrophs, phototrophs and plant pathogens. A study under controlled greenhouse conditions showed a reduction of the relative abundance of plant pathogenic protists was reduced by organic fertilizer amendments, while bacterivores and omnivores increased (Xiong et al., 2018). However, in a field study, plant parasites were favoured by organic fertilization (Harkes et al., 2019). Thus, currently a global and coherent picture of the responses of protistan plant pathogens to fertilization and other anthropogenic changes cannot be inferred.

Elucidating the diversity, dynamics and the environmental drivers of protistan plant pathogens is important for a sustainable land management, not only in agrosystems but also in natural ecosystems, because of their importance as drivers of patterns of plant diversity and productivity (Mitchell, 2003; Schnitzer et al., 2011). However, we are lacking baseline data on their occurrence in natural reservoirs, such as grasslands and forests (Stukenbrock and McDonald, 2008; van der Heijden et al., 2008). Cercozoa (Cavalier-Smith, 1998), a highly diverse phylum of c. 600 described species (Pawlowski et al., 2012), is a major protistan lineage in soil (Geisen et al., 2015; Grossmann et al., 2016; Lanzén et al., 2016; Ferreira de Araujo et al., 2018), comprizing a vast array of functional traits in morphologies, nutrition and locomotion modes (Burki and Keeling, 2014). Recently the phylum Endomyxa was separated from Cercozoa (Bass et al., 2018; Cavalier-Smith et al., 2018), and it is of particular interest for containing *inter alia* plant pathogens of functional and economic significance, including important transmitters of plant viruses and agents of root tumors (e.g. club root disease) (Neuhauser et al., 2014; Bass et al., 2019).

Much attention has been given to changes in biodiversity and ecosystem function following human disturbances, e.g. land use intensification in grasslands. A continuous, quantitative index (Land Use Intensity Index, LUI), integrating quantitative data on mowing, grazing and fertilization of grasslands (Blüthgen et al., 2012) has been widely applied in the framework of the German Biodiversity Exploratories. LUI had strong overall effects on ecosystem multifunctionality (Allan et al., 2015) and imperilled plant and animal community stability (Blüthgen et al., 2016). In particular, it was shown that plant species richness decreased along land use gradients (Egorov et al., 2014). With regard to soil microorganisms, LUI only weakly influenced bacterial communities (Kaiser et al., 2016), slightly increased ammonia oxidizing prokaryotes (Keil et al., 2011) and had no effect on protistan community assemblages as detected by T-RFLP (Glaser et al., 2015). However, it was demonstrated that specific environmental responses of protists remained undetectable when general eukaryotic PCR primers were used, but instead could be observed by the more thorough coverage of single lineages by taxon-specific primers (Lentendu et al., 2014)

Our aims were thus: (i) the assessment of the diversity of Cercozoa and Endomyxa, classified into trophic guilds, in grassland and forest along regional and land use intensification gradients; (ii) the identification of environmental factors driving the distribution of the cercozoan and endomyxan communities, with a special focus on plant pathogens. Our study took place in 150 grassland and 150 forest sites of the German Biodiversity Exploratories, sampled in 2011 and 2017 (600 soil samples in total). We tested the hypothesis that land use intensification would lead to shifts in the relative abundance of trophic guilds, increasing plant pathogens. Furthermore, we investigated to which degree grasslands and forests constituted natural reservoirs of the endomyxan plant parasites. Elucidating these patterns, both at regional and landscape scale, will provide new insights into the functional organization of the soil ecosystem and may give indications for a more sustainable land-use management.

## 2. Results

### 2.1 Sequencing results

We obtained c. 20 million reads per run (Table 1). Our primers were highly specific: the percentage of non-targeted OTUs accounted for only 8%. We obtained 2,101 genuine cercozoan OTUs from 600 grassland and forest soil samples representing nearly 18 million sequences. Nine samples were removed because the yield was not sufficient (<10,000 reads/sample), probably due to problems during the amplification, in 2011: SEG16, SEF31, SEF33, HEF07, HEF21, HEF23, HEF33; and in 2017: HEF07, SEF18. The remaining 591 samples yielded on average 29,817 reads/sample (min. 10,300, max 64,130, SD 9,784). The average number of OTUs per sample was 904 (maximum 1,365, minimum 254, SE 170.5). A database with the OTU abundance per sample, taxonomic assignment and estimated functional traits is provided (Table S1). Our sequencing and sampling efforts were sufficient, since the actual OTU richness was reached after only c. 235,000 sequences (Figure S2A) and at 65 samples (Figure S2B).

**Table 1.**
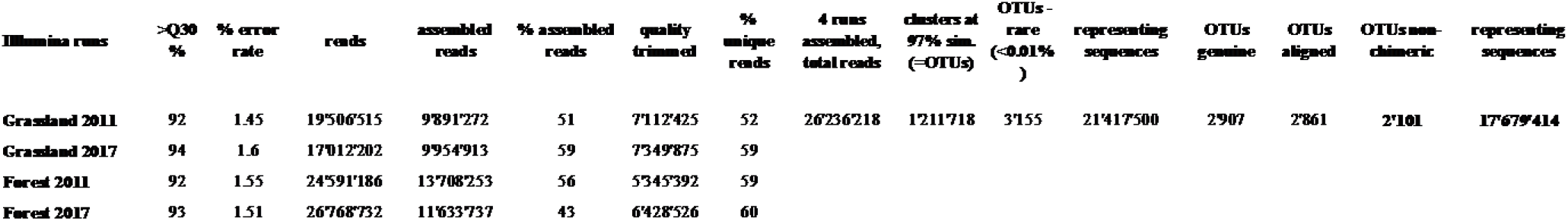
Quality estimates and error rate for each run; initial, assembled, quality-trimmed, and unique number of reads for each run. Number of reads retrieved at each step of the sequence processing for the 4 assembled runs.

### 2.2 Taxonomic and functional diversity of Cercozoa and Endomyxa

The 2,101 OTUs represented 329 unique Blast best hits. Only 28% of the OTUs were 97-100% similar to any known sequence (Figure S1). At a high taxonomic level, the majority of the OTUs could be assigned to the phylum Cercozoa (75%), the remaining to Endomyxa (25%) (Figure 1). Within Cercozoa, the class Sarcomonadea dominated (40%), composed of Glissomonadida and Cercomonadida, mostly composed of small flagellates and amoeboflagellates, respectively. Other main lineages were the testate amoeba with silica shells in Euglyphida (13%, Imbricatea) and those with mainly organic tests in Cryomonadida (12%, Thecofilosea). Endomyxa was composed of the plant parasitic Plasmodiophorida (14%, Phytomyxea) and amoeboid Vampyrellida (11%, Proteomyxidea). Naked cells (flagellates, ~19% and amoeboflagellates, ~32%, amoebae, ~10%) together constituted the predominant morphotypes, whereas testate cells (~22%) were less frequent. Among the latter, cells with organic/agglutinated test accounted for ~9% and those with a siliceous test for ~13-14%. Considering trophic positions, nearly 50% of the OTUs were bacterivores, but omnivores (~23%) and eukaryvores (~10 %) also contributed sizeable proportions, in both grasslands and forests. Compared to these predominant functional groups, the plant parasites (~5%) and hyperparasites of Oomycota (~0.8%) made up a much smaller proportion in grasslands and surprisingly, they were absent from forest soils. The overwhelmingly dominant locomotion mode was creeping/gliding in forest and grassland soils, suggesting a predominance of biofilm feeders (Figure 2).

**Figure 1.**
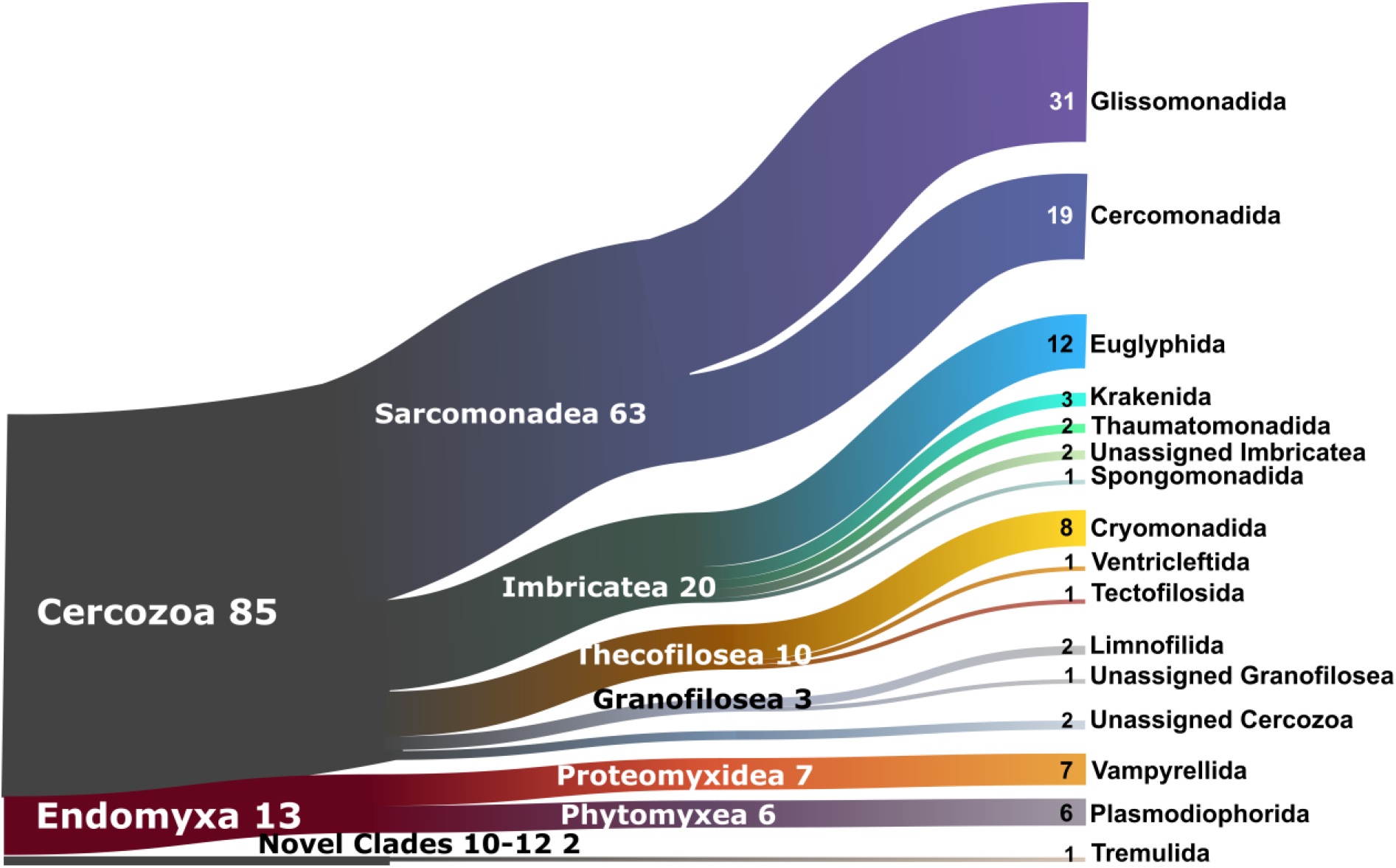
Sankey diagram showing the relative contribution of the cercozoan and endomyxan OTUs to the taxonomic diversity. Taxonomical assignment is based on the best hit by BLAST. From left to right, names refer to phylum (Cercozoa, Endomyxa), class (ending -ea) and orders (ending -ida). Numbers are percentages of OTUs abundance, taxa representing <1% are not shown.

**Figure 2.**
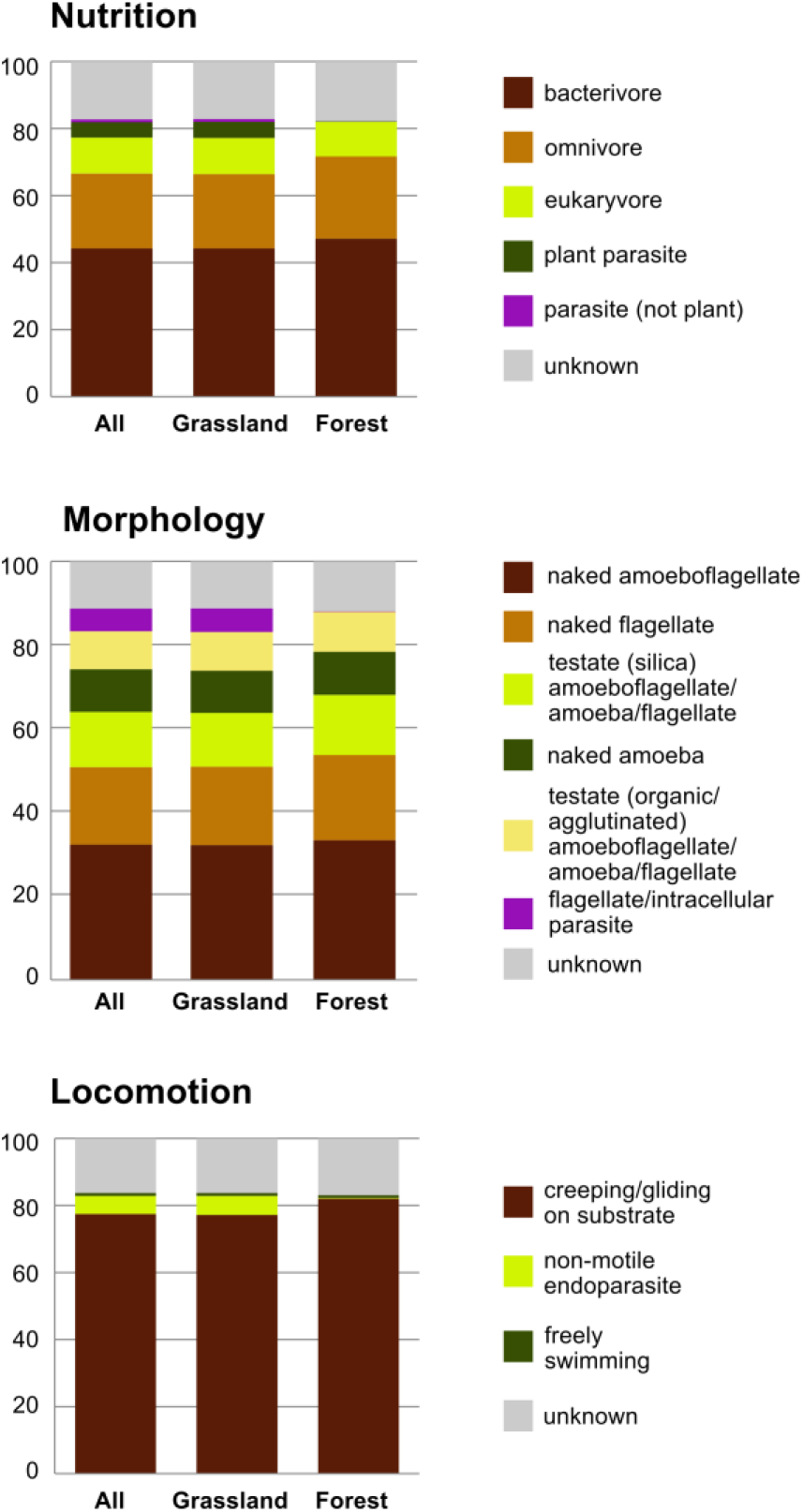
Histograms comparing the relative proportion of functional diversity classes of Cercozoa and Endomyxa (nutrition, morphology and locomotion) between the two ecosystems.

### 2.3 Alpha and beta diversity

A total of 2,004 and 1,808 OTUs were retrieved in grassland and forest sites, respectively. 1,712 OTUs (81%) were shared between grassland and forest, while only 293 and 97 OTUs were unique to grasslands and forests, respectively. Most OTUs were shared between regions, except two that were absent in Alb, one in Hainich, and three in Schorfheide (Table S1), and one present only in Schorfheide. Consequently, the influence of OTUs richness on protistan alpha diversity was low compared to that of relative abundances in the Shannon index. Thus, higher alpha diversity of Cercozoa and Endomyxa in forest compared to grassland were likely explained by a higher evenness (Figure S3). Despite the almost ubiquitous presence of protistan OTUs in soil samples of grasslands and forests, their communities dramatically differed between the two ecosystems. PCoA showed a strong arch effect indicating the overwhelming importance of the first component (x-axis), which separated forest and grassland sites, explaining 34% of the variance in Bray-Curtis distances (Figure 3). The second PCoA component explained 16% of variance and roughly reflected the regional differences along the south-north gradient, with the communities of Schorfheide standing out, especially in forests (Figure 3).

**Figure 3.**
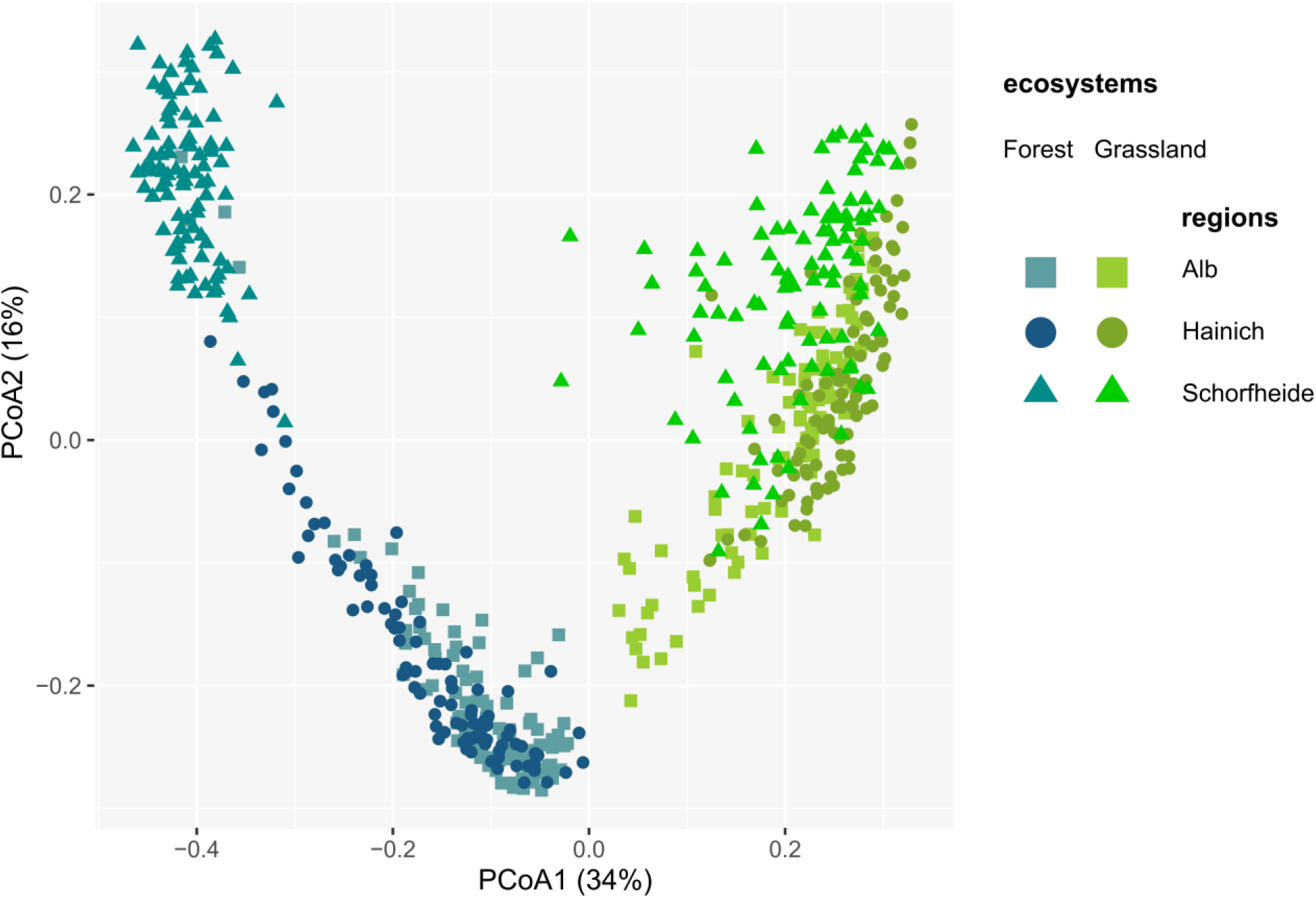
Principal Component Analysis of the Bray-Curtis dissimilarity indices between cercozoan and endomyxan OTUs, showing dramatically different communities between the two ecosystems and in a lesser extent between regions.

### 2.4 Major drivers of cercozoan and endomyxan communities turnover

Variance partitioning indicated that the ecosystem (forest vs grassland), the region (Alb, Hainich and Schorfheide) and the year of sampling (2011 vs 2017) together accounted for 42.7% (adjusted R^2^) of the total variation in beta diversity. Ecosystem explained 30.4% of the variation, followed by differences between regions (9.7%) and year of sampling (2.4%). Differences in soil properties and management practices always explained more variance of beta diversity in forests (R^2^ values 10.2-53.2) than in grasslands (R^2^ values 3.6-7.3) (Table S5). The most parsimonious models identified a strong influence of soil type and soil C/N ratio on protistan community composition for both grassland and forest (Table S5). In grassland, protistan communities were influenced by indicators of human disturbances, in particular differences in grassland management intensity (pastures, mowed meadows, meadows). This was also true for shifts of beta diversity within trophic guilds (except for bacterivores), as well as when both Cercozoa and Endomyxa were considered (all OTUs, Table S5). In addition, overall protistan beta diversity was influenced by land use intensification (according to the LUI index) and in particular by grassland fertilization. At the regional level, differences in LUI had a significant influence on protistan communities of Schorfheide and Hainich grasslands, with the additional effect of mowing intensity in Hainich, and the soil parameters clay content and C/N ratio in Schorfheide. Edaphic factors had a stronger influence on protistan communities in forest than in grassland. Most important factors were pH, clay content, C/N ratio, soil type and the identity of the dominant tree species (Table S5). This was true for all OTUs as well as for the individual trophic guilds. Bacterivores were further influenced by differences in the organic carbon content. At the regional level organic C was a stronger determinant than soil type. (Table S5).

Differences in the relative abundances of trophic guilds between regions were more pronounced in forests than in grasslands (Figure S4 & Figure S5). In grassland, the relative abundance of trophic guilds differed between regions, with bacterivores and omnivores less abundant in Hainich and Schorfheide, respectively (Figure S4A). Soil types significantly influenced the relative abundances of bacterivores and omnivores; bacterivores were most abundant in Leptosol and less abundant in Stagnosol; omnivores were most abundant in Leptosol and Cambisol and less abundant in Histosol and Gleysol (Figure S4B). The relative abundance of plant parasites increased with a more intense grassland management (from pastures to mowed meadows and meadows) (Figure S4C), with a higher land use intensity (Figure S4D) and decreased with a higher C/N ratio (Figure S4E). Omnivores and eukaryvores were more abundant at low LUI (Figure S4D) and at high C/N ratio (Figure S4E)

In forest, the patterns shown by bacterivores and eukaryvores were opposite: factors favouring one functional group impaired the other - with omnivores showing a somewhat intermediate trend (Figure S5). Bacterivores were markedly more abundant in Schorfheide, while eukaryvores showed the opposite trend, while the relative abundances of omnivores only slightly differed between the three regions (Figure S5A). All trophic guilds were differentially affected by soil type (Figure S5B). The identity of the dominant tree species had a strong effect on bacterivores, which were less abundant under beech and spruce than pine and oak - while the opposite was true for eukaryvores (Figure S5C). Edaphic factors had a much stronger influence on trophic guilds in forests than in grasslands. The relative abundance of bacterivores was highest at low organic carbon content, while the eukaryvores showed the opposite trend; omnivores only slightly increased at a high organic C content (Figure S5D). Bacterivores increased and eukaryvores decreased at low pH (3-4), while omnivores slightly increased at high pH (Figure S5E). Increasing Organic C, pH, clay content and decreased C/N ratio all depleted the bacterivores and increased the eukaryvores (Figure S5D, E, F, and G).

## 3 Discussion

Our study was based on a thorough sampling of protists across three regions spanning a South-North gradient in Germany. Sequencing depth reached saturation (Figure S2), a precondition to detect detailed responses of each functional group to the ecological processes involved in shaping their distribution. The high percentage of OTUs not closely matching any published sequence (Figure S1) indicated a significant hidden species richness not yet taxonomically recorded or sequenced. Consistently with previous studies on Cercozoa and Endomyxa, a high alpha-diversity and a low beta-diversity were found (Lentendu et al., 2018; Fiore-Donno et al., 2019). Our data not only confirmed the low endemicity of Cercozoa, but due to our thorough sampling, high sequencing depth and the use of taxon-specific primers, we showed that almost all OTUs were shared between ecosystems and regions. This implies that community assembly of these protists is not limited by dispersal over countrywide distances. As a corollary, the remarkable differences in beta diversity then must reflect differences in how taxonomic and trophic guilds respond to environmental selection (Fiore-Donno et al., 2019).

### 3.1 Phytomyxea are absent from forests

One remarkable exception to the ubiquitous occurrence of our taxa was the endomyxan plant parasites (Phytomyxea) which, apart of three OTUs, were not represented in the forest ecosystem (Figure 2 & Table S5). To our knowledge, this striking pattern was not observed to date, although Ferreira de Araujo et al. (2018) noted that grass-dominated ecosystems hosted more plant parasites than tree-dominated ones. The absence of Phytomyxea in temperate forests is corroborated by a recent metatranscriptomics study on leaf litter of 18 Biodiversity Exploratories forest sites (Voss et al., 2019), but they were detected in low abundance in tropical forest soils (Mahé et al., 2017). The dominant endomyxan OTUs in grasslands were identified as *Polymyxa graminis*, *Spongospora nasturtii* and *Woronina pythii* (Table S2). The first two are plant parasites associated with grasses and Brassicaceae, respectively. Since both grasses and Brassicaceae are present in forests, although to a lesser extent than in grasslands, the remarkable absence of Phytomyxea from the 300 forest soils cannot be attributed to the lack of suitable hosts. Instead, it might be hypothesized that the establishment of the parasites depends on a threshold density of their host plants, or that the forest soil microbial community has a suppressing effect on Phytomyxea. Both hypotheses deserve further attention due their potential applied value for sustainable agriculture. *Woronina pythii* is an endomyxan hyperparasite of plant-parasitic oomycetes, the latter belonging to the supergroup Stramenopiles. Our study confirmed that *Woronina* is widespread and relatively abundant in soils (Neuhauser et al., 2014; Fiore-Donno et al., 2019), suggesting a potential use as a biological pest control.

### 3.2 Responses to human-induced disturbances in grasslands

The most parsimonious model suggested that both relative abundance and beta diversity of endomyxan plant parasites in grasslands were influenced by management type and the region-dependent soil types (Table S5), but were unresponsive to other edaphic factors. Their relative abundances increased with more intense grassland management (from pasture to meadows) (Figure S4C), with a higher LUI index (Figure S4D) and corresponding shifts towards higher soil nutrient contents (lower C/N, 7-10) (Figure S4E). In the Biodiversity Exploratories, the LUI index mainly depended on N fertilization and mowing (Blüthgen et al., 2012). Thus nitrogen-rich, intensively managed meadows constitute an important natural reservoir for these plant parasites. Beta diversity of cercozoan and endomyxan OTUs was influenced by anthropogenic changes such as land use intensification (LUI index), fertilization, and differences in the soil C/N ratio. This confirms a recent study where protists were identified as the most susceptible component of the soil microbiome to the application of nitrogen fertilizers in agricultural soils (Zhao et al., 2019). Mowing intensity had a pronounced negative effect on plant species richness in Hainich (Socher et al., 2013). Accordingly, the significant influence of mowing intensity on protistan beta diversity in Hainich (Table S5) was likely driven by shifts in plant species composition.

### 3.3 Opposite responses of trophic guilds in forest

Bacterivores and eukaryvores showed opposite responses in their regional abundances to different environmental parameters in forest (Figure S5 & Figure 4), while omnivores did not respond so markedly, probably because they combine bacterivory and eukaryvory. Since bacterivores and eukaryvores rely on entirely different food sources, their opposite responses may reflect shifts in the availability of bacteria and microbial eukaryotes linked to specific forest soil conditions. For example, the increase of bacterivores and decrease of eukaryvores in Schorfheide could be explained by the lower biomass of soil fungi in this region (Birkhofer et al., 2012). Bacterivores and eukaryvores were differentially influenced by three edaphic factors, i.e. pH, texture and organic carbon content (Figure 4), the very same major drivers of bacterial community structure at national and continental scales (Bahram et al., 2018; Karimi et al., 2018; Plassart et al., 2019).

**Figure 4.**
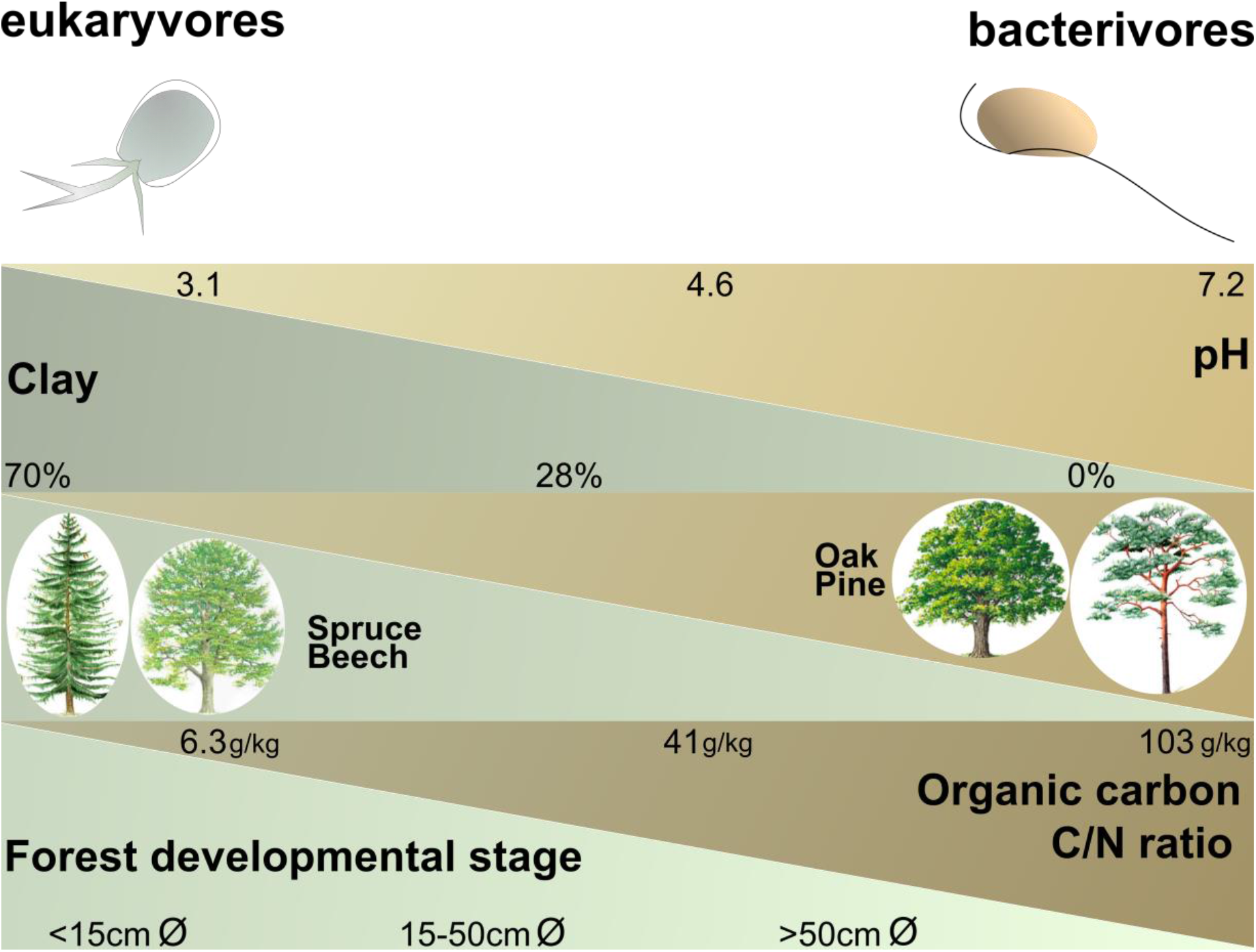
Schematic illustration showing the positive or negative influences of the most influencial ecological and edaphic parameters in forest (according to the models in Table S5) and the relative abundances of the cercozoan bacterivores and eukaryvores. Minimum, average and maximum values are given for each parameter. Forest developmental stage was estimated by the mean of the diameter of 100 trunks of well-developed trees of the dominant species. The parameters are not co-correlated: e.g. bacterivores are favoured by soils with a higher pH or by soils with a higher organic carbon content (not necessarily by both).

### 3.4 Soil type and C/N ratio as major structuring forces of protistan beta diversity

Generally, soil type appeared as a major determinant of differences in cercozoan and endomyxan beta diversity between the three regions, confirming previous observations using microbial phospho-lipid fatty acids (PLFAs) (Herold et al., 2014). Soil types also influenced beta diversity within and relative proportions between different trophic guilds, including bacterivores, omnivores and eukaryvores (Figure 4 & Figure S5). The importance of soil type is corroborated by significant effects of soil physico-chemical parameters (soil pH, clay content, soil organic matter content) on the regional scale (Table S5 & Figure S5). Soil types, with their specific physical and chemical properties, are considered to be strong environmental filters for the assembly of microbial communities, sometimes more influential than plant species (Garbeva et al., 2004; Harkes et al., 2019). Importantly, soil types may contribute to explain regional differences in microbial community composition.

C/N ratio was identified as a robust structuring factor of belowground biota across biomes (Fierer et al., 2009), and it had been shown to decrease microbial biomass (Dequiedt et al., 2011; Serna-Chavez et al., 2013). In our study sites, C/N ratios were lower in grassland (7-15) than in forest (11-26) (Figure S4 & Figure S5). Unexpectedly, while eukaryvores followed the expected pattern of microbial biomass, we found bacterivores to be favoured by a high C/N ratio in forest, although such substrates would limit the biomass of bacteria.

Regional differences among the Biodiversity Exploratories are marked: the three regions differ substantially in diversity of macro organisms, bacteria and environmental conditions (Allan et al., 2015; Kaiser et al., 2016). Different assemblages of protists reflected differences between the three regions (Glaser et al., 2015; Fiore-Donno et al., 2016), although not in the study of Venter et al. (2017), something the authors attribute to under sampling. In our study, the regional differences were more pronounced in forest than in grassland (Figure 3, Figure S4A & Figure S5A), with Schorfheide standing out. In particular, the marked differences of abundances of bacterivores and eukaryvores in Schorfheide were mirrored in the types of soil and trees only present in this region (Figure S5).

### 3.5 Strong differences in protistan community composition between grassland and forest

The main driver shaping the cercozoan and endomyxan communities was the ecosystem, with striking differences between grassland and forest in terms of alpha- and beta-diversity, as shown by Shannon index, evenness, principal component analysis and variance partitioning (Figure S3 & Figure 3). Similar to our results, a higher richness in forests compared to grasslands and/or agricultural fields and shrublands was reported for protists (albeit not for Cercozoa) (Ferreira de Araujo et al., 2018) and eukaryotes in general (Zhao et al., 2018). A similar trend was observed in the Biodiversity Exploratories for bacteria (Nacke et al., 2001; Foesel et al., 2014; Kaiser et al., 2016) and arbuscular mycorrhizal and saprophytic fungi (Birkhofer et al., 2012).

In our settings, the year of sampling only explained a negligible proportion of the variance, probably because in both years sampling took place in April. Thus, the seasonal dependence of protistan communities collected across one year of sampling in one of the investigated grassland plots in Alb (Fiore-Donno et al., 2019), could not be observed. This suggests that the seasonal dynamics tend to be similar every year.

## 4. Conclusion

Our intensive study on temperate grassland and forest was based on an established protocol for metabarcoding of two important protistan lineages, classified into ecologically meaningful guilds, which allowed distinguish important trends in community dynamics and biogeographies. Such trends could be masked in studies considering only high-ranking groups of microorganisms, since opposite responses of different trophic guilds - as shown in this study - could add up to a neutral response. Our data identified management practices favouring plant parasites in grassland, mostly linked to N fertilization. Further investigations on co-occurrences of protistan trophic guilds with bacteria and fungi may further reveal consumer-resource relationships in soil food webs of grassland and forest ecosystems.

## 5. Materials and Methods

### 5.1 Study sites, soil sampling and DNA extraction

Our study took place in three German Biodiversity Exploratories, i.e. the Biosphere Reserve Schorfheide-Chorin in the State of Brandenburg, the National Park Hainich and its surroundings in the State of Thuringia and the Biosphere Reserve Schwäbische Alb in the State of Baden-Württemberg (Fischer et al., 2010). Each exploratory comprises 50 grassland sites from extensive pastures to highly fertilized meadows and 50 differently managed forest sites. Each site contains a study plot of 20 × 20 m. From all study plots, 300 soil samples were collected in a coordinated joint sampling campaign within 14 days in April 2011 and a second one in April 2017. From each plot 14 soil cores of 8.3 cm diameter were taken every 3 m along two transects of 20 m each, oriented North-South and East-West, employing a soil corer. The surface layer (0-10 cm) was collected, after removing plants, pebbles and conspicuous roots. Soil cores from each plot were sieved (2 mm mesh size), mixed, homogenised and immediately frozen for further analysis. Soil DNA was extracted from 400 mg of soil, 3- to 6-times, using the DNeasy PowerSoil Kit (Qiagen GmbH, Hilden, Germany) following the manufacturer’s protocol, to obtain a sufficient amount to be shared between research groups of the Biodiversity Exploratories.

### 5.2 Primer and barcode design, amplification and sequencing

We modified the primers specific for Cercozoa from Fiore-Donno et al. (2018) to better match the parasitic lineages Phytomyxea and Proteomyxidea in Endomyxa. The first PCR was conducted with a mixture of the forward primers S615F_Cerco + S615F_Phyt, 50% each (all primers 5’-3’) GTTAAAAAGCTCGTAGTTG and GTTAAAARGCTCGTAGTCG and reverse S963R_Phyt CAACTTTCGTTCTTGATYAAA; the second PCR was conducted with barcoded primers (Table S2), forward S615F_Cer GTTAAAARGCTCGTAGTYG and reverse S947R_Cer AAGARGAYATCCTTGGTG. The barcodes consisted in eight-nucleotide-long sequences appended to the 5’-ends of both the forward and the reverse primers, because tagging only one primer leads to extensive mistagging (Esling et al., 2015). To design the barcodes, we first used barcrawl (Frank, 2009) to obtain a list of barcodes with a balanced nucleotide content (no homopolymers), not folding on themselves and to themselves an the attached primer (no “hairpin”), not forming heteroduplexes with the corresponding primer and having at least 3 bases differences between them. In addition, using custom R scripts, we selected from the previous list only the barcodes that did not match the consensus of the reference alignment flanking the primer region and without cross-dimerization between each combination of primer+barcodes. We designed 18 barcoded versions for the forward and the reverse primers (Table S2), allowing for 324 possible combinations to label samples of which only 150 were used, since it is advisable to leave a proportion of unused combinations to decrease mistagging (Esling et al., 2015). Barcoded primers were specifically ordered for NGS application to Microsynth (Wolfurt, Austria).

For the amplification, we incorporated 1 μl of 1:10 soil DNA template for the first PCR round and 1 μl of the resulting amplicons as a template for a following semi-nested PCR. We employed the following final concentrations: GreenTaq polymerase (Fermentas, Canada) 0.01units, buffer 1x, dNTPs 0.2 mM and primers 1 μM. The thermal programme consisted of an initial denaturation step at 95°C for 2 min, 24 cycles at 95°C for 30 s, 50°C for 30 s, 72°C for 30 s; and a final elongation step at 72°C for 5 min. The number of PCR cycles was kept at 24 since chimera formation arises dramatically after 25 cycles (Michu et al., 2010). All PCRs were conducted twice to reduce the possible artificial dominance of few amplicons by PCR competition (2 × 10 μl for the first and 2 × 27 μl for the second PCR), and the two amplicons were pooled after the second PCR.

The amplicons were checked by electrophoresis and 25 μl of each were purified and normalized using SequalPrep Normalization Plate Kit (Invitrogen GmbH, Karlsruhe, Germany) to obtain a concentration of 1–2 ng/μl per sample, and the 150 samples were pooled. We conducted 4 Illumina runs, for each year (2011 and 2017) and for grassland or forest. During the library preparation, amplicons were end-repaired, small fragments were removed, 3’ ends were adenylated, and Illumina adapters and sequencing primers were ligated (TruSeqDNA PCR-Free, Illumina Inc., San Diego, CA, USA). The library was quantified by qPCR, performed following the manufacturer’s instructions (KAPA SYBR^®^ FAST qPCR Kit, Kapabiosystems, Wilmington, MA, USA) on a CFX96 Real Time System (Bio-Rad, Hercules, CA, USA). Sequencing was performed with a MiSeq v3 Reagent kit of 300 cycles (on a MiSeq Desktop Sequencer (Illumina Inc., San Diego, CA, USA) at the University of Geneva (Switzerland), Department of Genetics and Evolution.

### 5.3 Sequences processing

Paired reads were assembled using MOTHUR v.39.5 (Schloss et al., 2009) (which was also used in the following steps) allowing no differences in the primers and the barcodes, no ambiguities and removing assembled sequences <300bp and with an overlap <100bp (Table 1). The quality check and removal/cutting of low-quality reads were conducted with the default parameters. Reads were sorted into samples via detection of the barcodes (Table S2) and sequences renamed with the sample, year and “G” for grassland or “F” for forest. After clustering identical sequences together, all reads were assembled in one file. Sequences were clustered into Operational Taxonomic Units (OTUs) using VSEARCH (Rognes et al., 2016) with abundance-based greedy clustering (agc) and a similarity threshold of 97%. Clusters represented by less than 0.01% of the reads were removed as likely to represent amplification or sequencing noise (Fiore-Donno et al., 2018). Using BLAST+ (Camacho et al., 2008) with an e-value of 1^e-50^ and keeping only the best hit, sequences were identified in the PR2 database (Guillou et al., 2013) and non cercozoan sequences were removed. Sequences were aligned with the provided template (Fiore-Donno et al., 2018) with a penalty for opening gaps of −5. Chimeras were identified using UCHIME (Edgar et al., 2011) as implemented in mothur and chimeric sequences removed (Table 1). The results are shown as a table with the OTUs abundance/site, their taxonomic assignment according to the best hit by Blast and their functional assignment according to Dumack et al. (2019). The relative abundance of each cercozoan taxonomic level was illustrated using Sankey diagram generator V1.2 (http://sankey-diagram-generator.acquireprocure.com/, last accessed Jan. 2019) and refined with the open-source vector graphic editor Inkscape (https://inkscape.org/en/, last accessed Jan. 2019).

### 5.4 Statistical analyses

All statistical analyses were carried out within the R environment (R v. 3.5.1) (R Development Core Team, 2014), on the OTU abundance/sites table (Table S1), and the environmental parameters (Table S3 & Table S4), the latter normalized by the K-nearest neighbours. Unless otherwise specified, community analyses were performed with the package vegan (Oksanen et al., 2013). **Alpha diversity**: To evaluate if more sampling and sequencing effort would have revealed more OTU richness, we carried out an analysis based on OTUs accumulation curves, function *specaccum*, method rarefaction and 1,000 random permutations; species richness was extrapolated using the function *specpool*. Alpha diversity estimates were based on relative abundances of OTUs (function *decostand*, method “total”); Shannon diversity and Pielou’s evenness were obtained with the function *diversity*. **Beta-diversity**: Variation partitioning (function *varpart* applied to the Hellinger-transformed OTUs dataset) was applied to assess the amount of explained beta diversity by the factors region, year of collection (2011, 2017) and or ecosystem (grassland vs forest). Beta diversity between regions and ecosystems was inferred by Principal Coordinate Analysis (PCoA, function *cmdscale*), using Bray-Curtis dissimilarities (function *vegdist*, method “bray”) on the relative abundances of OTUs, then plotted with the package ggplot2. A distance-based redundancy analysis (dbRDA) on Bray-Curtis dissimilarities (function *dbrda*) was used to investigate the effect of environmental factors on the beta-diversity of cercozoan and endomyxan communities. Parsimonious models were selected by the function *ordistep* with default parameters based on 999 permutations, and only significant results were shown in dbRDA. All available parameters were tested for co-correlation according to variance inflation factors (function *vif.cca*). Among the soil parameters measured in the framework of the Biodiversity Exploratories (Table S3 & Table S4), soil carbon (total, organic and inorganic) and nitrogen (total N) were all co-correlated; based on slightly higher significance in the models organic carbon was kept. Among the three co-correlated descriptors of soil texture, soil clay content was chosen over sand and silt contents.

Because the relative abundance is an important component of the beta diversity, the effects of environmental parameters on the **relative abundances** of the trophic guilds were tested with general linear models (function *glm*, core package). They were then subjected to the general linear hypothesis test (function *glht*, package multcomp) with Tukey’s test for multiple comparisons of means and a heteroskedasticity-consistent covariance matrix estimation (function *vcovHC*, package sandwich).

## 6. Data accessibility

Raw sequences have been deposited in Sequence Read Archive (NCBI) Bioproject PRJNA513166, SRA10697391-4 and the 2,101 OTUs (representative sequences) under GenBank accession numbers MN322900-MN325000.

## 7. Author contributions

MB and AMFD conceived the PATHOGEN project. MB administrated the project. AMFD conducted the amplifications, Illumina sequencing and bioinformatics pipeline. AMFD and TRH performed statistics. AMFD, TRH and MB interpreted the data. AMFD wrote the manuscript. All co-authors participated in the revisions and approved the final version of the manuscript

## 8. Acknowledgments

This work was funded by the German Research Foundation (DFG) in the frame of the priority program 1374 Biodiversity Exploratories [grant number BO 1907/18–1] (PATHOGEN). We are very grateful to Linhui Jiang and Cristopher Kahlich for invaluable help in the lab. We thank Graham Jones, Scotland for writing the R scripts selecting barcodes. At the University of Geneva (CH), we thank Jan Pawlowski and Emanuela Reo and we are liable to the Swiss National Science Foundation Grant 316030 150817 funding the MiSeq instrument. We thank the managers of the Exploratories, Swen Renner, Kirsten Reichel-Jung, Kerstin Wiesner, Katrin Lorenzen, Martin Gorke, Miriam Teuscher, and all former managers for their work in maintaining the plot and project infrastructure; Simone Pfeiffer and Christiane Fischer for giving support through the central office, Jens Nieschulze and Michael Owonibi for managing the central data base, and Markus Fischer, Eduard Linsenmair, Dominik Hessenmüller, Daniel Prati, Ingo Schöning, François Buscot, Ernst-Detlef Schulze, Wolfgang W. Weisser and the late Elisabeth Kalko for their role in setting up the Biodiversity Exploratories project. Field work permits were issued by the responsible state environmental offices of Baden-Württemberg, Thüringen, and Brandenburg.

## Supplementary material

### Supplementary tables

**Table S1**. Database of the abundance of each cercozoan and endomyxan OTU per sample. The taxonomic assignment (super group, class, order, family, genus and species) is provided according to the best hit by BLAST (PR2 database), with the % of similarity. Functional traits (morphology, nutrition and locomotion modes) were estimated to the genus level following Dumack et al. (2019).

**Table S2**. Combinations of barcodes used in this study, with the corresponding soil samples. Code for sample names: AE=Alb, HE=Hainich, SE=Schorfheide. G=grassland sites - to be replaced by F in forest sites (the same barcodes were applied since the grassland and forest samples were amplified and sequenced separately).

**Table S3**. Environmental parameters from the 150 grassland study sites and two years of collection.

**Table S4**. Environmental parameters from the 150 forest study sites and two years of collection.

**Table S5**. Most parsimonious models (dbRDA), with their respective R^2^ adjusted and F values, and the F values of the factors selected by each model. Significance values shown as symbol (see footnote).

### Supplementary figures

**Figure S1**. Similarities of the cercozoan and endomyxan OTUs with known sequences. OTUs are classified according to their percentage of similarity to the next kin by BLAST. The horizontal bar length is proportional to the number of OTUs in each rank. Shaded area=OTUs with a similarity ≥ 97%.

**Figure S2**. Description of the cercozoan and endomyxan diversity. A. Rarefaction curve describing the observed number of OTUs as a function of the sequencing effort; saturation was reached with c. 235,000 sequences. B. Species accumulation curve describing the sampling effort; saturation was reached with 65 samples.

**Figure S3**. Boxplots and table of the alpha diversity of the cercozoan and endomyxan OTUs estimated with the Shannon and evenness indices, for all sites and for sites binned by region and ecotype. Red letters: a change from “a” to “b”, or “c” indicates a significant difference (multiple comparison of means, Tukey’s test); two or three letters (e.g. “ab” or “abc”) indicate non-significant differences between plots sharing those letters. Red dots indicate the means (black lines the median). In the table, the highest means are in bold.

**Figure S4**. Boxplots of the variation of the relative abundances of the four main nutrition modes of Cercozoa and Endomyxa in grassland, coloured according to region. **A.** by region; **B.** by soil type, only Cambisoil is found in the three regions; **C.** by grassland management; **D.** by land use intensity (LUI) index, transformed into a categorical variable according to quantiles. **E**. by C/N ratio, transformed as in D. The y-scale varies between graphs. Red letters: a change from “a” to “b”, or “c” indicates a significant difference (multiple comparison of means, Tukey’s test); two or three letters (e.g. “ab” or “abc”) indicate non-significant differences between plots sharing those letters. Red lines indicate the mean.

**Figure S5**. Boxplots of the variation of the relative abundances of the three main nutrition modes of Cercozoa in forest (nearly no plant parasites were found in forest), coloured according to region. **A.** by region; **B.** by soil type, only Cambisoil is found in two regions; **C.** by main tree species, Latin names: beech = *Fagus sylvatica*, spruce = *Picea abies*, pine = *Pinus sylvestris*, oak = *Quercus petrea* & *Q. robur*, pine and oak growing only in Schorfheide; **D**. by organic carbon (g/kg soil) levels; **E**. by pH levels; **F**. by levels of percentage of soil clay content. **G**. by levels of C/N ratio. The y-scale varies between graphs. Red letters: a change from “a” to “b”, or “c” indicates a significant difference (multiple comparison of means, Tukey’s test); two or three letters (e.g. “ab” or “abc”) indicate non-significant differences between boxes sharing those letters. Red lines indicate the mean.

