## Supplementary material for "Contrasting responses of protistan plant parasites and phagotrophs to ecosystems, land management and soil properties": Table S2

**Table S1**. Combinantion of barcodes used to identify soil samples. A stands for Schwäbische Alb; H for Hainich and S for Schorfheide-Chorin. G stands for grassland, F for forest.

| Primer | GTTAAAARGCTCGTAGTYG | AAGARGAYATCCTTGGTG | Grassland sites | Forest sites |
|---|---|---|---|---|
| barcode | GTGAACTC | GTCAGTAT | AEG001 | AEF001 |
| barcode | GTGAACTC | TACGCTAT | AEG002 | AEF002 |
| barcode | GTGAACTC | TTAGGAAC | AEG003 | AEF003 |
| barcode | GTGAACTC | GTAACATG | AEG004 | AEF004 |
| barcode | GTGAACTC | GACATATC | AEG005 | AEF005 |
| barcode | GTGAACTC | TACTGTAG | AEG006 | AEF006 |
| barcode | GTGAACTC | TCTCTCAG | AEG007 | AEF007 |
| barcode | GTGAACTC | ACGATCAG | AEG008 | AEF008 |
| barcode | GCGTAATC | TATCAGTC | AEG009 | AEF009 |
| barcode | GCGTAATC | GCTTCAAT | AEG010 | AEF010 |
| barcode | GCGTAATC | AATCAGGT | AEG011 | AEF011 |
| barcode | GCGTAATC | ACAATGTG | AEG012 | AEF012 |
| barcode | GCGTAATC | AATTGGTC | AEG013 | AEF013 |
| barcode | GCGTAATC | AAGCTACT | AEG014 | AEF014 |
| barcode | GCGTAATC | ATTCTCGG | AEG015 | AEF015 |
| barcode | GCGTAATC | GTGTCAAC | AEG016 | AEF016 |
| barcode | ATGTGACC | TACGCTAT | AEG017 | AEF017 |
| barcode | ATGTGACC | TTAGGAAC | AEG018 | AEF018 |
| barcode | ATGTGACC | GTAACATG | AEG019 | AEF019 |
| barcode | ATGTGACC | GACATATC | AEG020 | AEF020 |
| barcode | ATGTGACC | TACTGTAG | AEG021 | AEF021 |
| barcode | ATGTGACC | TCTCTCAG | AEG022 | AEF022 |
| barcode | ATGTGACC | CGTTCAAG | AEG023 | AEF023 |
| barcode | ATGTGACC | ACGATCAG | AEG024 | AEF024 |
| barcode | ATACGCAC | TATCAGTC | AEG025 | AEF025 |
| barcode | ATACGCAC | GCTTCAAT | AEG026 | AEF026 |
| barcode | ATACGCAC | AATCAGGT | AEG027 | AEF027 |
| barcode | ATACGCAC | CCATTATG | AEG028 | AEF028 |
| barcode | ATACGCAC | ACAATGTG | AEG029 | AEF029 |
| barcode | ATACGCAC | AATTGGTC | AEG030 | AEF030 |
| barcode | ATACGCAC | ATTCTCGG | AEG031 | AEF031 |
| barcode | ATACGCAC | GTGTCAAC | AEG032 | AEF032 |
| barcode | ATCTGGAC | TACGCTAT | AEG033 | AEF033 |
| barcode | ATCTGGAC | TTAGGAAC | AEG034 | AEF034 |
| barcode | ATCTGGAC | GTAACATG | AEG035 | AEF035 |
| barcode | ATCTGGAC | GACATATC | AEG036 | AEF036 |
| barcode | ATCTGGAC | TACTGTAG | AEG037 | AEF037 |
| barcode | ATCTGGAC | TCTCTCAG | AEG038 | AEF038 |
| barcode | ATCTGGAC | CGTTCAAG | AEG039 | AEF039 |
| barcode | ATCTGGAC | ACGATCAG | AEG040 | AEF040 |
| barcode | GTCAATCC | GCTTCAAT | AEG041 | AEF041 |
| barcode | GTCAATCC | AATCAGGT | AEG042 | AEF042 |
| barcode | GTCAATCC | CCATTATG | AEG043 | AEF043 |
| barcode | GTCAATCC | ACAATGTG | AEG044 | AEF044 |
| barcode | GTCAATCC | AATTGGTC | AEG045 | AEF045 |
| barcode | GTCAATCC | AAGCTACT | AEG046 | AEF046 |
| barcode | GTCAATCC | ATTCTCGG | AEG047 | AEF047 |

| | | | | |
|---|---|---|---|---|
| barcode | GTCAATCC | GTGTCAAC | AEG048 | AEF048 |
| barcode | GTCAATCC | TATCAGTC | AEG049 | AEF049 |
| barcode | ATGTGACC | GTCAGTAT | AEG050 | AEF050 |
| barcode | GACCATAC | TACGCTAT | HEG001 | HEF001 |
| barcode | GACCATAC | TTAGGAAC | HEG002 | HEF002 |
| barcode | GACCATAC | GTAACATG | HEG003 | HEF003 |
| barcode | GACCATAC | GACATATC | HEG004 | HEF004 |
| barcode | GACCATAC | TACTGTAG | HEG005 | HEF005 |
| barcode | GACCATAC | TCTCTCAG | HEG006 | HEF006 |
| barcode | GACCATAC | CGTTCAAG | HEG007 | HEF007 |
| barcode | GACCATAC | ACGATCAG | HEG008 | HEF008 |
| barcode | GAATACTC | TATCAGTC | HEG009 | HEF009 |
| barcode | GAATACTC | GCTTCAAT | HEG010 | HEF010 |
| barcode | GAATACTC | AATCAGGT | HEG011 | HEF011 |
| barcode | GAATACTC | CCATTATG | HEG012 | HEF012 |
| barcode | GAATACTC | ACAATGTG | HEG013 | HEF013 |
| barcode | GAATACTC | AAGCTACT | HEG014 | HEF014 |
| barcode | GAATACTC | ATTCTCGG | HEG015 | HEF015 |
| barcode | GAATACTC | GTGTCAAC | HEG016 | HEF016 |
| barcode | GTTATAGC | GTCAGTAT | HEG017 | HEF017 |
| barcode | GTTATAGC | TTAGGAAC | HEG018 | HEF018 |
| barcode | GTTATAGC | GTAACATG | HEG019 | HEF019 |
| barcode | GTTATAGC | GACATATC | HEG020 | HEF020 |
| barcode | GTTATAGC | TACTGTAG | HEG021 | HEF021 |
| barcode | GTTATAGC | TCTCTCAG | HEG022 | HEF022 |
| barcode | GTTATAGC | CGTTCAAG | HEG023 | HEF023 |
| barcode | GTTATAGC | ACGATCAG | HEG024 | HEF024 |
| barcode | TAGTTACC | TATCAGTC | HEG025 | HEF025 |
| barcode | TAGTTACC | GCTTCAAT | HEG026 | HEF026 |
| barcode | TAGTTACC | CCATTATG | HEG027 | HEF027 |
| barcode | TAGTTACC | ACAATGTG | HEG028 | HEF028 |
| barcode | TAGTTACC | AATTGGTC | HEG029 | HEF029 |
| barcode | TAGTTACC | AAGCTACT | HEG030 | HEF030 |
| barcode | TAGTTACC | ATTCTCGG | HEG031 | HEF031 |
| barcode | TAGTTACC | GTGTCAAC | HEG032 | HEF032 |
| barcode | AACTTAGC | TACGCTAT | HEG033 | HEF033 |
| barcode | AACTTAGC | TTAGGAAC | HEG034 | HEF034 |
| barcode | AACTTAGC | GTAACATG | HEG035 | HEF035 |
| barcode | AACTTAGC | GACATATC | HEG036 | HEF036 |
| barcode | AACTTAGC | TACTGTAG | HEG037 | HEF037 |
| barcode | AACTTAGC | TCTCTCAG | HEG038 | HEF038 |
| barcode | AACTTAGC | CGTTCAAG | HEG039 | HEF039 |
| barcode | AACTTAGC | ACGATCAG | HEG040 | HEF040 |
| barcode | TATCTAGC | GCTTCAAT | HEG041 | HEF041 |
| barcode | TATCTAGC | AATCAGGT | HEG042 | HEF042 |
| barcode | TATCTAGC | CCATTATG | HEG043 | HEF043 |
| barcode | TATCTAGC | ACAATGTG | HEG044 | HEF044 |
| barcode | TATCTAGC | AATTGGTC | HEG045 | HEF045 |
| barcode | TATCTAGC | AAGCTACT | HEG046 | HEF046 |
| barcode | TATCTAGC | ATTCTCGG | HEG047 | HEF047 |
| barcode | TATCTAGC | GTGTCAAC | HEG048 | HEF048 |
| barcode | AACTTAGC | GTCAGTAT | HEG049 | HEF049 |

| | | | | |
|---|---|---|---|---|
| barcode | GACCATAC | GTCAGTAT | HEG050 | HEF050 |
| barcode | TGATACAC | GTCAGTAT | SEG001 | SEF001 |
| barcode | TGATACAC | TACGCTAT | SEG002 | SEF002 |
| barcode | TGATACAC | TTAGGAAC | SEG003 | SEF003 |
| barcode | TGATACAC | GTAACATG | SEG004 | SEF004 |
| barcode | TGATACAC | GACATATC | SEG005 | SEF005 |
| barcode | TGATACAC | TACTGTAG | SEG006 | SEF006 |
| barcode | TGATACAC | TCTCTCAG | SEG007 | SEF007 |
| barcode | TGATACAC | CGTTCAAG | SEG008 | SEF008 |
| barcode | GTAGATAC | GCTTCAAT | SEG009 | SEF009 |
| barcode | GTAGATAC | AATCAGGT | SEG010 | SEF010 |
| barcode | GTAGATAC | CCATTATG | SEG011 | SEF011 |
| barcode | GTAGATAC | ACAATGTG | SEG012 | SEF012 |
| barcode | GTAGATAC | AATTGGTC | SEG013 | SEF013 |
| barcode | GTAGATAC | AAGCTACT | SEG014 | SEF014 |
| barcode | GTAGATAC | ATTCTCGG | SEG015 | SEF015 |
| barcode | GTAGATAC | GTGTCAAC | SEG016 | SEF016 |
| barcode | AGTTCATC | GTCAGTAT | SEG017 | SEF017 |
| barcode | AGTTCATC | TACGCTAT | SEG018 | SEF018 |
| barcode | AGTTCATC | TTAGGAAC | SEG019 | SEF019 |
| barcode | AGTTCATC | GTAACATG | SEG020 | SEF020 |
| barcode | AGTTCATC | GACATATC | SEG021 | SEF021 |
| barcode | AGTTCATC | TACTGTAG | SEG022 | SEF022 |
| barcode | AGTTCATC | CGTTCAAG | SEG023 | SEF023 |
| barcode | AGTTCATC | ACGATCAG | SEG024 | SEF024 |
| barcode | TTAGAGTC | GCTTCAAT | SEG025 | SEF025 |
| barcode | TTAGAGTC | AATCAGGT | SEG026 | SEF026 |
| barcode | TTAGAGTC | CCATTATG | SEG027 | SEF027 |
| barcode | TTAGAGTC | ACAATGTG | SEG028 | SEF028 |
| barcode | TTAGAGTC | AATTGGTC | SEG029 | SEF029 |
| barcode | TTAGAGTC | AAGCTACT | SEG030 | SEF030 |
| barcode | TTAGAGTC | ATTCTCGG | SEG031 | SEF031 |
| barcode | TTAGAGTC | GTGTCAAC | SEG032 | SEF032 |
| barcode | TATATGCC | TACGCTAT | SEG033 | SEF033 |
| barcode | TATATGCC | TTAGGAAC | SEG034 | SEF034 |
| barcode | TATATGCC | GTAACATG | SEG035 | SEF035 |
| barcode | TATATGCC | GACATATC | SEG036 | SEF036 |
| barcode | TATATGCC | TACTGTAG | SEG037 | SEF037 |
| barcode | TATATGCC | TCTCTCAG | SEG038 | SEF038 |
| barcode | TATATGCC | CGTTCAAG | SEG039 | SEF039 |
| barcode | TATATGCC | ACGATCAG | SEG040 | SEF040 |
| barcode | AATTGGTC | TATCAGTC | SEG041 | SEF041 |
| barcode | AATTGGTC | GCTTCAAT | SEG042 | SEF042 |
| barcode | AATTGGTC | AATCAGGT | SEG043 | SEF043 |
| barcode | AATTGGTC | CCATTATG | SEG044 | SEF044 |
| barcode | AATTGGTC | ACAATGTG | SEG045 | SEF045 |
| barcode | AATTGGTC | AATTGGTC | SEG046 | SEF046 |
| barcode | AATTGGTC | AAGCTACT | SEG047 | SEF047 |
| barcode | AATTGGTC | ATTCTCGG | SEG048 | SEF048 |
| barcode | TATATGCC | GTCAGTAT | SEG049 | SEF049 |
| barcode | TTAGAGTC | TATCAGTC | SEG050 | SEF050 |
| barcode | GTAGATAC | TATCAGTC | MOCK | MOCK |
