## Supplementary material for "Contrasting responses of protistan plant parasites and phagotrophs to ecosystems, land management and soil properties": Table S3

### Table S3. Environmental parameters from the 150 grassland study sites and two years of collection.

| Reference dataset at https://www.bexis.uni-jena.de/ | 14447 22246 | 14446 23846 | 14446 23846 | 14446 23846 | 14446 23846 | 14446 23846 | 14686 | 14686 | 14686 | 19266 | | | | 10580 | 10580 |
|---|---|---|---|---|---|---|---|---|---|---|---|---|---|---|---|
| Sites (AE=Alb, HE=Hainich, SE=Schorfheide, 11/17:year of collection | pH | Total_C g/kg soil | Inorganic_C g/kg | Organic_C g/kg | Total_N g/kg | CN_ ratio | Clay | Silt | Sand | LUI | Grazing[1] | Mowing[2] | Fertilization (kg N/ha) | soil_type | Grassland_management |
| AEG001_11 | 6.53 | 85.95 | 0.96 | 84.98 | 8.71 | 9.76 | 643 | 324 | 33 | 1.87 | 0 | 2 | 92.75 | Leptosol | meadow |
| AEG002_11 | 6.75 | 72.92 | 1.87 | 71.04 | 7.49 | 9.48 | 512 | 411 | 77 | 3.21 | 0 | 3 | 239 | Leptosol | meadow |
| AEG003_11 | 6.22 | 59.81 | 0.28 | 59.53 | 5.83 | 10.21 | 672 | 298 | 30 | 1.94 | 0 | 2 | 64.6 | Leptosol | meadow |
| AEG004_11 | 5.08 | 44.53 | 0.22 | 44.31 | 4.83 | 9.17 | 488 | 427 | 85 | 1.56 | 47.4 | 1 | 35 | Leptosol | mowed_meadow |
| AEG005_11 | 6.14 | 90.77 | 0.57 | 90.19 | 8.72 | 10.34 | 365 | 518 | 117 | 1.52 | 37 | 1 | 35 | Leptosol | mowed_meadow |
| AEG006_11 | 5.87 | 83.14 | 0.4 | 82.74 | 8.21 | 10.07 | 588 | 380 | 32 | 1.78 | 128 | 1 | 35 | Leptosol | mowed_meadow |
| AEG007_11 | 7.11 | 86.32 | 33.51 | 52.8 | 4.64 | 11.37 | 385 | 427 | 188 | 0.63 | 42.9 | 0 | 0 | Leptosol | pasture |
| AEG008_11 | 6.52 | 86.63 | 0.48 | 86.14 | 7.68 | 11.21 | 673 | 319 | 8 | 1.4 | 119 | 1 | 0 | Leptosol | pasture |
| AEG009_11 | 6.7 | 71.21 | 0.93 | 70.27 | 5.9 | 11.91 | 556 | 417 | 27 | 0.9 | 85.5 | 0 | 0 | Leptosol | pasture |
| AEG010_11 | 6.08 | 94.16 | 0.45 | 93.71 | 9.1 | 10.29 | 647 | 327 | 26 | 0.91 | 0 | 1 | 0 | Leptosol | meadow |
| AEG011_11 | 5.49 | 62.91 | 0.31 | 62.61 | 6.02 | 10.39 | 643 | 327 | 30 | 2.05 | 0 | 3 | 101.76 | Leptosol | meadow |
| AEG012_11 | 6.57 | 69.15 | 1.62 | 67.54 | 6.59 | 10.25 | 436 | 512 | 52 | 1.87 | 0 | 2 | 200.55 | Leptosol | meadow |
| AEG013_11 | 5.95 | 68.88 | 0.44 | 68.43 | 6.58 | 10.39 | 643 | 311 | 46 | 1.87 | 0 | 2 | 200.55 | Leptosol | meadow |
| AEG014_11 | 6.71 | 62.33 | 1.06 | 61.27 | 6.07 | 10.09 | 521 | 453 | 26 | 2.14 | 0 | 2 | 89.6 | Leptosol | meadow |
| AEG015_11 | 5.68 | 63.9 | 0.28 | 63.62 | 6.51 | 9.77 | 642 | 340 | 18 | 2.63 | 0 | 3 | 135.66 | Leptosol | meadow |
| AEG016_11 | 6.61 | 60.7 | 0.72 | 59.98 | 5.93 | 10.11 | 528 | 449 | 23 | 1.57 | 83.7 | 2 | 0 | Leptosol | mowed_meadow |
| AEG017_11 | 6.86 | 67.23 | 2.55 | 64.68 | 6.54 | 9.89 | 618 | 365 | 17 | 2.08 | 0 | 2 | 80.75 | Leptosol | meadow |
| AEG018_11 | 6.87 | 66.09 | 3.94 | 62.14 | 6.31 | 9.84 | 649 | 332 | 19 | 2.11 | 0 | 2 | 84.6 | Leptosol | meadow |
| AEG019_11 | 5.85 | 78.05 | 0.38 | 77.67 | 7.64 | 10.17 | 637 | 327 | 36 | 1.61 | 65.1 | 1 | 35 | Leptosol | mowed_meadow |
| AEG020_11 | 6.62 | 81.25 | 0.84 | 80.41 | 8.64 | 9.31 | 597 | 334 | 69 | 1.81 | 350 | 0 | 0 | Leptosol | pasture |
| AEG021_11 | 5.99 | 67.75 | 0.34 | 67.42 | 6.76 | 9.98 | 587 | 391 | 22 | 3.72 | 1226 | 1 | 7.95 | Leptosol | pasture |
| AEG022_11 | 5.88 | 53.56 | 0 | 53.56 | 4.8 | 11.16 | 708 | 270 | 22 | 1.08 | 35.1 | 1 | 106.48 | Leptosol | meadow |
| AEG023_11 | 6.77 | 83.43 | 3.16 | 80.28 | 8.17 | 9.83 | 566 | 387 | 47 | 0.91 | 0 | 1 | 0 | Leptosol | meadow |
| AEG024_11 | 6.08 | 80.7 | 0.47 | 80.23 | 7.78 | 10.31 | 638 | 329 | 33 | 1.84 | 164 | 2 | 47.3 | Leptosol | mowed_meadow |
| AEG025_11 | 6.97 | 99.27 | 17.13 | 82.13 | 7.29 | 11.27 | 463 | 406 | 131 | 0.76 | 61.9 | 0 | 47.3 | Leptosol | pasture |
| AEG026_11 | 6.66 | 70.02 | 0.6 | 69.42 | 5.73 | 12.12 | 679 | 241 | 80 | 1.81 | 350 | 0 | 0 | Leptosol | pasture |
| AEG027_11 | 6.15 | 89.96 | 0.49 | 89.47 | 7.69 | 11.63 | 576 | 410 | 14 | 1.2 | 153 | 0 | 0 | Leptosol | pasture |
| AEG028_11 | 6.08 | 54.34 | 0.31 | 54.04 | 4.83 | 11.19 | 599 | 385 | 16 | 0.83 | 73.1 | 0 | 0 | Leptosol | pasture |
| AEG029_11 | 5.77 | 53.67 | 0.3 | 53.37 | 5.17 | 10.32 | 566 | 343 | 91 | 1.36 | 66 | 1 | 19.8 | Cambisol | mowed_meadow |
| AEG030_11 | 6.65 | 65.91 | 0.95 | 64.96 | 6.68 | 9.73 | 556 | 375 | 69 | 1.79 | 254 | 1 | 94.6 | Leptosol | mowed_meadow |
| AEG031_11 | 6.55 | 70.21 | 0.58 | 69.63 | 6.83 | 10.18 | 595 | 381 | 24 | 1.95 | 318 | 1 | 0 | Leptosol | mowed_meadow |
| AEG032_11 | 5.3 | 42.05 | 0.26 | 41.79 | 3.93 | 10.64 | 461 | 466 | 73 | 0.75 | 59.6 | 0 | 0 | Leptosol | pasture |
| AEG033_11 | 5.98 | 49.21 | 0.28 | 48.93 | 4.61 | 10.61 | 597 | 388 | 15 | 1.32 | 186 | 0 | 0 | Leptosol | pasture |
| AEG034_11 | 6.17 | 46.23 | 0.33 | 45.9 | 4.72 | 9.72 | 423 | 554 | 23 | 1.02 | 111 | 0 | 0 | Leptosol | pasture |
| AEG035_11 | 5.67 | 39.54 | 0 | 39.54 | 4.03 | 9.81 | 446 | 503 | 51 | 1.83 | 0 | 2 | 45 | Cambisol | meadow |
| AEG036_11 | 6.02 | 53.74 | 0.42 | 53.33 | 5.14 | 10.38 | 588 | 213 | 199 | 1.84 | 0 | 2 | 101.76 | Cambisol | meadow |
| AEG037_11 | 6.1 | 63.43 | 0.43 | 63 | 6.31 | 9.99 | 566 | 334 | 100 | 1.68 | 0 | 2 | 35 | Cambisol | meadow |
| AEG038_11 | 5.51 | 49.45 | 0 | 49.45 | 5.13 | 9.64 | 451 | 468 | 81 | 1.29 | 0 | 2 | 0 | Cambisol | meadow |
| AEG039_11 | 5.67 | 66.41 | 0.32 | 66.09 | 6.28 | 10.52 | 463 | 493 | 44 | 1.68 | 0 | 2 | 35 | Cambisol | meadow |
| AEG040_11 | 6.71 | 69.95 | 1.6 | 68.34 | 7.02 | 9.74 | 698 | 269 | 33 | 1.64 | 0 | 3 | 47.3 | Cambisol | meadow |
| AEG041_11 | 6.15 | 44.56 | 0.28 | 44.28 | 4.31 | 10.26 | 541 | 407 | 52 | 2.25 | 26 | 2 | 117.84 | Cambisol | meadow |
| AEG042_11 | 6.97 | 96.45 | 13.31 | 83.13 | 9.23 | 9 | 159 | 757 | 84 | 2.14 | 85 | 2 | 64.6 | Cambisol | mowed_meadow |
| AEG043_11 | 6.81 | 75.63 | 3.06 | 72.57 | 7.47 | 9.71 | 613 | 349 | 38 | 1.73 | 150 | 0 | 48.45 | Cambisol | mowed_meadow |
| AEG044_11 | 7.05 | 78.49 | 14.47 | 64.01 | 6.65 | 9.63 | 152 | 735 | 113 | 2.18 | 504 | 0 | 0 | Cambisol | pasture |
| AEG045_11 | 5.19 | 55.07 | 0.2 | 54.86 | 5.72 | 9.6 | 529 | 417 | 54 | 1.29 | 0 | 2 | 0 | Cambisol | meadow |
| AEG046_11 | 5.68 | 71.91 | 0.23 | 71.68 | 7.25 | 9.88 | 659 | 287 | 54 | 1.66 | 293 | 0 | 0 | Cambisol | pasture |
| AEG047_11 | 7.19 | 104.4 | 45.38 | 59.05 | 5.47 | 10.8 | 146 | 704 | 150 | 0.81 | 69.3 | 0 | 0 | Cambisol | pasture |
| AEG048_11 | 7.3 | 97.98 | 54.58 | 43.39 | 3.38 | 12.84 | 295 | 565 | 140 | 0.75 | 59.1 | 0 | 0 | Cambisol | pasture |
| AEG049_11 | 6.16 | 50.14 | 0.27 | 49.87 | 4.81 | 10.36 | 451 | 522 | 27 | 1.05 | 117 | 0 | 0 | Cambisol | pasture |

| | | | | | | | | | | | | | | | |
|---|---|---|---|---|---|---|---|---|---|---|---|---|---|---|---|
| AEG050_11 | 5.93 | 68.98 | 0.31 | 68.68 | 6.76 | 10.15 | 589 | 382 | 29 | 1.57 | 0 | 2 | 189.2 | Cambisol | meadow |
| HEG001_11 | 6.65 | 54.78 | 0.76 | 54.02 | 5.46 | 9.89 | 502 | 447 | 50 | 2.42 | 0 | 2 | 150 | Cambisol | meadow |
| HEG002_11 | 7.2 | 39.23 | 4.15 | 35.08 | 3.55 | 9.88 | 552 | 396 | 52 | 2.63 | 0 | 3 | 150 | Vertisol | meadow |
| HEG003_11 | 7.26 | 39.17 | 5.88 | 33.29 | 3.4 | 9.81 | 544 | 399 | 57 | 2.63 | 0 | 3 | 150 | Vertisol | meadow |
| HEG004_11 | 6.65 | 65.32 | 0.6 | 64.72 | 6.21 | 10.42 | 511 | 414 | 75 | 2.18 | 180 | 1 | 81 | Stagnosol | mowed_meadow |
| HEG005_11 | 7.09 | 48.4 | 4.8 | 43.6 | 4.44 | 9.82 | 454 | 497 | 49 | 2.61 | 149 | 3 | 90 | Stagnosol | mowed_meadow |
| HEG006_11 | 5.96 | 20.77 | 0 | 20.77 | 2.01 | 10.33 | 257 | 698 | 45 | 2.32 | 107 | 1 | 132 | Stagnosol | mowed_meadow |
| HEG007_11 | 6.99 | 57.9 | 2.41 | 55.48 | 5.7 | 9.73 | 536 | 434 | 30 | 0.58 | 36 | 0 | 0 | Stagnosol | pasture |
| HEG008_11 | 7.17 | 60.63 | 3.59 | 57.04 | 5.78 | 9.86 | 494 | 452 | 56 | 0.58 | 36 | 0 | 0 | Stagnosol | pasture |
| HEG009_11 | 6.92 | 42.26 | 1.14 | 41.11 | 3.58 | 11.49 | 387 | 537 | 76 | 0.75 | 59.5 | 0 | 0 | Stagnosol | pasture |
| HEG010_11 | 6.4 | 44.03 | 0.3 | 43.74 | 4.01 | 10.9 | 436 | 532 | 30 | 1.59 | 99.3 | 1 | 170.28 | Vertisol | meadow |
| HEG011_11 | 7.25 | 65.07 | 8.44 | 56.64 | 5.45 | 10.4 | 552 | 405 | 43 | 1.46 | 57.4 | 1 | 170.28 | Stagnosol | meadow |
| HEG012_11 | 6.98 | 78.38 | 3.49 | 74.89 | 7.86 | 9.53 | 497 | 448 | 55 | 3.42 | 1143 | 0 | 40 | Stagnosol | mowed_meadow |
| HEG013_11 | 7.17 | 38.26 | 2.82 | 35.44 | 3.57 | 9.93 | 394 | 536 | 70 | 1.58 | 40 | 2 | 0 | Stagnosol | mowed_meadow |
| HEG014_11 | 6.28 | 39.21 | 0.33 | 38.88 | 3.61 | 10.75 | 450 | 491 | 60 | 2 | 19.8 | 2 | 116.76 | Stagnosol | mowed_meadow |
| HEG015_11 | 7.1 | 55.11 | 1.79 | 53.32 | 5.24 | 10.16 | 528 | 435 | 38 | 1.82 | 38.7 | 1 | 75 | Stagnosol | mowed_meadow |
| HEG016_11 | 6.61 | 64.89 | 0.85 | 64.04 | 6.39 | 10.02 | 555 | 405 | 39 | 1.24 | 164 | 0 | 0 | Stagnosol | pasture |
| HEG017_11 | 6.99 | 54.11 | 0.94 | 53.17 | 4.96 | 10.71 | 546 | 419 | 32 | 0.77 | 63.9 | 0 | 0 | Stagnosol | pasture |
| HEG018_11 | 7.32 | 66.61 | 18.47 | 48.14 | 4.1 | 11.74 | 457 | 446 | 97 | 0.81 | 70.6 | 0 | 0 | Vertisol | pasture |
| HEG019_11 | 6.59 | 62.68 | 0.5 | 62.18 | 5.71 | 10.89 | 485 | 427 | 87 | 0.77 | 63.9 | 0 | 0 | Stagnosol | pasture |
| HEG020_11 | 5.4 | 27.23 | 0 | 27.23 | 2.34 | 11.66 | 239 | 661 | 102 | 0.69 | 51.4 | 0 | 0 | Stagnosol | pasture |
| HEG021_11 | 7.33 | 42.07 | 10.33 | 31.73 | 3.02 | 10.5 | 311 | 631 | 58 | 0.61 | 39.6 | 0 | 0 | Stagnosol | pasture |
| HEG022_11 | 6.92 | 50.12 | 2.24 | 47.88 | 4.94 | 9.7 | 446 | 467 | 87 | 1.52 | 18.7 | 2 | 0 | Cambisol | mowed_meadow |
| HEG023_11 | 7.2 | 50.17 | 3.09 | 47.08 | 4.75 | 9.9 | 588 | 375 | 37 | 1.28 | 62.3 | 1 | 0 | Stagnosol | mowed_meadow |
| HEG024_11 | 6.63 | 48.98 | 0.81 | 48.17 | 5.08 | 9.48 | 545 | 404 | 52 | 1.45 | 110 | 1 | 0 | Stagnosol | mowed_meadow |
| HEG025_11 | 7.26 | 59.54 | 4.34 | 55.2 | 5.6 | 9.87 | 423 | 531 | 46 | 1.77 | 334 | 0 | 0 | Cambisol | pasture |
| HEG026_11 | 7.29 | 56.14 | 8.66 | 47.48 | 4.7 | 10.1 | 481 | 458 | 61 | 1.46 | 0 | 2 | 0 | Cambisol | meadow |
| HEG027_11 | 7.29 | 48.76 | 4.28 | 44.49 | 4.26 | 10.43 | 492 | 472 | 36 | 1.6 | 0 | 1 | 60 | Cambisol | meadow |
| HEG028_11 | 7.22 | 38.19 | 6.12 | 32.08 | 3.26 | 9.85 | 536 | 409 | 55 | 1.76 | 0 | 1 | 81 | Cambisol | mowed_meadow |
| HEG029_11 | 7.12 | 30.56 | 1.02 | 29.54 | 2.87 | 10.28 | 469 | 501 | 30 | 1.76 | 0 | 1 | 81 | Cambisol | mowed_meadow |
| HEG030_11 | 7.11 | 40.15 | 4.64 | 35.51 | 3.67 | 9.67 | 400 | 548 | 52 | 2.22 | 0 | 3 | 70 | Cambisol | mowed_meadow |
| HEG031_11 | 7.15 | 46.29 | 5.26 | 41.03 | 4.09 | 10.02 | 237 | 726 | 36 | 2.25 | 104 | 1 | 121 | Cambisol | mowed_meadow |
| HEG032_11 | 5.53 | 40.19 | 0.22 | 39.97 | 3.79 | 10.54 | 340 | 640 | 17 | 2.01 | 158 | 1 | 60 | Cambisol | mowed_meadow |
| HEG033_11 | 5.02 | 40.08 | 0 | 40.08 | 3.84 | 10.45 | 353 | 618 | 29 | 2.1 | 140 | 1 | 81 | Cambisol | pasture |
| HEG034_11 | 7.06 | 35.28 | 1.33 | 33.95 | 3.4 | 9.97 | 448 | 469 | 83 | 2.32 | 107 | 1 | 132 | Cambisol | mowed_meadow |
| HEG035_11 | 6.97 | 50.8 | 5.05 | 45.74 | 4.79 | 9.56 | 307 | 580 | 113 | 2.32 | 107 | 1 | 132 | Cambisol | mowed_meadow |
| HEG036_11 | 7.25 | 63.76 | 10.7 | 53.06 | 5.32 | 9.97 | 61 | 871 | 71 | 2.04 | 69.6 | 1 | 98 | Cambisol | mowed_meadow |
| HEG037_11 | 7.41 | 66.94 | 14.32 | 52.61 | 5.39 | 9.75 | 60 | 841 | 99 | 2.1 | 69.6 | 1 | 108 | Cambisol | mowed_meadow |
| HEG038_11 | 7.29 | 33.77 | 5.44 | 28.33 | 2.84 | 9.98 | 310 | 622 | 68 | 1.67 | 185 | 1 | 0 | Cambisol | pasture |
| HEG039_11 | 6.46 | 38.87 | 0.43 | 38.44 | 3.9 | 9.85 | 351 | 537 | 110 | 1.24 | 165 | 0 | 0 | Cambisol | pasture |
| HEG040_11 | 6.73 | 61.48 | 1.07 | 60.41 | 5.99 | 10.08 | 275 | 648 | 79 | 2.06 | 454 | 0 | 0 | Cambisol | pasture |
| HEG041_11 | 7.21 | 41.98 | 2.17 | 39.81 | 3.52 | 11.29 | 518 | 417 | 65 | 0.69 | 51.4 | 0 | 0 | Cambisol | pasture |
| HEG042_11 | 7.15 | 44 | 2.19 | 41.81 | 3.83 | 10.91 | 435 | 520 | 46 | 0.77 | 63.9 | 0 | 0 | Cambisol | pasture |
| HEG043_11 | 7.16 | 46.85 | 1.44 | 45.41 | 4.07 | 11.16 | 376 | 589 | 38 | 1.42 | 101 | 1 | 0 | Cambisol | pasture |
| HEG044_11 | 7.15 | 84.38 | 4.81 | 79.57 | 7.7 | 10.32 | 79 | 853 | 69 | 0.76 | 61.4 | 0 | 0 | Cambisol | pasture |
| HEG045_11 | 6.73 | 44.99 | 0.55 | 44.44 | 4.27 | 10.39 | 492 | 468 | 43 | 0.85 | 76.8 | 0 | 0 | Cambisol | pasture |
| HEG046_11 | 7.45 | 47.28 | 21.44 | 25.83 | 2.48 | 10.4 | 395 | 531 | 74 | 0.87 | 80.6 | 0 | 0 | Cambisol | pasture |
| HEG047_11 | 7.09 | 58.25 | 3.16 | 55.09 | 5.57 | 9.89 | 483 | 477 | 40 | 1.86 | 141 | 2 | 0 | Cambisol | mowed_meadow |
| HEG048_11 | 6.8 | 42.18 | 0.54 | 41.64 | 4.03 | 10.33 | 465 | 488 | 50 | 1.75 | 215 | 1 | 0 | Cambisol | mowed_meadow |
| HEG049_11 | 6.63 | 47.17 | 0.67 | 46.5 | 4.65 | 10 | 431 | 500 | 71 | 1.51 | 16.3 | 2 | 0 | Cambisol | mowed_meadow |
| HEG050_11 | 6.85 | 45.95 | 0.59 | 45.36 | 4.42 | 10.27 | 646 | 335 | 23 | 1.15 | 26.4 | 1 | 0 | Cambisol | mowed_meadow |
| SEG001_11 | 7.24 | 218.5 | 14.93 | 203.5 | 19.1 | 10.65 | 233 | 466 | 301 | 3.1 | 0 | 3 | 50 | Histosol | meadow |
| SEG002_11 | 7.32 | 171.2 | 42.76 | 128.4 | 12.7 | 10.1 | 198 | 527 | 275 | 2.82 | 88.9 | 1 | 50 | Histosol | meadow |
| SEG003_11 | 7.42 | 135.5 | 45.7 | 89.82 | 8.89 | 10.1 | 89 | 600 | 311 | 2.92 | 0 | 2 | 50 | Histosol | meadow |
| SEG004_11 | 7.35 | 210.5 | 69.4 | 141.1 | 13.9 | 10.15 | 163 | 686 | 151 | 1.03 | 0 | 1 | 0 | Histosol | mowed_meadow |
| SEG005_11 | 7.42 | 181.2 | 66.56 | 114.7 | 11.51 | 9.96 | 162 | 575 | 263 | 1.46 | 0 | 2 | 0 | Gleysol | mowed_meadow |
| SEG006_11 | 5.54 | 284.5 | 3.95 | 280.6 | 22.51 | 12.46 | 248 | 284 | 468 | 1.49 | 413 | 0 | 0 | Histosol | mowed_meadow |
| SEG007_11 | 7.3 | 174.5 | 59.42 | 115 | 11.04 | 10.42 | 157 | 648 | 195 | 1.59 | 472 | 0 | 0 | Histosol | pasture |
| SEG008_11 | 7.34 | 157.2 | 78.55 | 78.71 | 7.34 | 10.71 | 110 | 713 | 177 | 1.03 | 0 | 1 | 0 | Gleysol | pasture |
| SEG009_11 | 6.56 | 233.6 | 4.11 | 229.4 | 18.2 | 12.6 | 171 | 143 | 686 | 0.94 | 163 | 0 | 0 | Histosol | pasture |
| SEG010_11 | 7.4 | 205.4 | 58.16 | 147.3 | 14.21 | 10.36 | 183 | 516 | 301 | 2.92 | 0 | 2 | 50 | Histosol | meadow |
| SEG011_11 | 7.43 | 183.8 | 71.32 | 112.5 | 11.48 | 9.8 | 137 | 552 | 311 | 2.92 | 0 | 2 | 50 | Gleysol | meadow |
| SEG012_11 | 7.37 | 144.4 | 38.59 | 105.8 | 10.65 | 9.93 | 138 | 548 | 314 | 3.1 | 0 | 3 | 50 | Histosol | meadow |
| SEG013_11 | 7.13 | 35.92 | 2.65 | 33.27 | 3.37 | 9.89 | 41 | 324 | 635 | 1.46 | 0 | 2 | 0 | Cambisol | meadow |

| | | | | | | | | | | | | | | | |
|---|---|---|---|---|---|---|---|---|---|---|---|---|---|---|---|
| SEG014_11 | 7.35 | 172.3 | 68.55 | 103.8 | 10.45 | 9.94 | 169 | 625 | 206 | 0.82 | 124 | 0 | 0 | Gleysol | pasture |
| SEG015_11 | 7.37 | 193.4 | 57.21 | 136.2 | 14.09 | 9.67 | 191 | 593 | 216 | 1.46 | 0 | 2 | 0 | Histosol | mowed_meadow |
| SEG016_11 | 7.4 | 200.9 | 60.07 | 140.8 | 13.97 | 10.07 | 178 | 655 | 167 | 1.03 | 0 | 1 | 0 | Gleysol | mowed_meadow |
| SEG017_11 | 5.03 | 355.8 | 4.11 | 351.7 | 27.77 | 12.66 | 374 | 348 | 278 | 1.15 | 243 | 0 | 0 | Histosol | mowed_meadow |
| SEG018_11 | 5.14 | 14.87 | 0 | 14.87 | 1.21 | 12.3 | 65 | 89 | 846 | 1.46 | 0 | 2 | 0 | Luvisol | meadow |
| SEG019_11 | 7.33 | 101.4 | 25.68 | 75.74 | 7.51 | 10.09 | 87 | 553 | 360 | 1.2 | 71 | 1 | 0 | Gleysol | pasture |
| SEG020_11 | 6.59 | 276 | 6.49 | 269.5 | 21.54 | 12.51 | 307 | 304 | 389 | 1.4 | 165 | 1 | 0 | Histosol | pasture |
| SEG021_11 | 5.35 | 283 | 2.71 | 280.3 | 21.84 | 12.83 | 278 | 310 | 412 | 1.21 | 271 | 0 | 0 | Gleysol | pasture |
| SEG022_11 | 7.28 | 107.9 | 44.61 | 63.35 | 6.53 | 9.7 | 93 | 636 | 271 | 2.29 | 974 | 0 | 0 | Gleysol | pasture |
| SEG023_11 | 5.2 | 296.6 | 3.68 | 292.9 | 20.71 | 14.14 | 373 | 461 | 166 | 2.5 | 0 | 2 | 32.3 | Histosol | meadow |
| SEG024_11 | 7.07 | 232.1 | 28.81 | 203.3 | 20.16 | 10.08 | 283 | 504 | 213 | 1.03 | 0 | 1 | 0 | Histosol | meadow |
| SEG025_11 | 6.31 | 202.3 | 4.56 | 197.8 | 16.21 | 12.19 | 176 | 206 | 618 | 1.46 | 0 | 2 | 0 | Histosol | meadow |
| SEG026_11 | 7.14 | 305 | 7.94 | 297.1 | 24.17 | 12.29 | 427 | 353 | 220 | 2.5 | 0 | 2 | 32.3 | Histosol | meadow |
| SEG027_11 | 5.76 | 370.8 | 11.29 | 359.5 | 24.69 | 14.56 | 416 | 417 | 167 | 1.03 | 0 | 1 | 0 | Histosol | meadow |
| SEG028_11 | 7.3 | 202.3 | 56.98 | 145.3 | 13.63 | 10.66 | 175 | 651 | 174 | 1.03 | 0 | 1 | 0 | Histosol | meadow |
| SEG029_11 | 7.39 | 181 | 49.45 | 131.6 | 12.94 | 10.16 | 195 | 694 | 110 | 1.03 | 0 | 1 | 0 | Histosol | meadow |
| SEG030_11 | 7.09 | 31.96 | 2.56 | 29.4 | 2.79 | 10.52 | 146 | 343 | 511 | 1.46 | 0 | 2 | 0 | Albeluvisol | meadow |
| SEG031_11 | 6.24 | 28.79 | 0.36 | 28.43 | 2.64 | 10.77 | 182 | 281 | 537 | 1.46 | 0 | 2 | 0 | Cambisol | meadow |
| SEG032_11 | 5.91 | 24.18 | 0 | 24.18 | 2.3 | 10.5 | 190 | 234 | 576 | 1.46 | 0 | 2 | 0 | Luvisol | meadow |
| SEG033_11 | 5.71 | 18.22 | 0 | 18.22 | 1.75 | 10.41 | 118 | 265 | 617 | 1.32 | 325 | 0 | 0 | Albeluvisol | mowed_meadow |
| SEG034_11 | 5.58 | 18.41 | 0 | 18.41 | 1.74 | 10.59 | 138 | 273 | 589 | 1.58 | 151 | 0 | 85.14 | Albeluvisol | mowed_meadow |
| SEG035_11 | 5.97 | 22.83 | 0 | 22.83 | 2.15 | 10.62 | 200 | 238 | 562 | 1.71 | 35.7 | 1 | 85.14 | Luvisol | mowed_meadow |
| SEG036_11 | 6.2 | 24.58 | 0.29 | 24.28 | 2.41 | 10.09 | 139 | 301 | 560 | 1.36 | 344 | 0 | 0 | Albeluvisol | pasture |
| SEG037_11 | 4.58 | 12.22 | 0 | 12.22 | 1.16 | 10.51 | 93 | 99 | 808 | 1.38 | 156 | 1 | 0 | Albeluvisol | pasture |
| SEG038_11 | 5.17 | 22.82 | 0 | 22.82 | 2.18 | 10.46 | 89 | 72 | 838 | 1.57 | 259 | 1 | 0 | Cambisol | mowed_meadow |
| SEG039_11 | 7.07 | 22.29 | 5.42 | 16.87 | 1.63 | 10.34 | 225 | 237 | 538 | 1.42 | 176 | 1 | 0 | Cambisol | pasture |
| SEG040_11 | 5.76 | 21.3 | 0 | 21.3 | 2.03 | 10.49 | 98 | 192 | 710 | 1.74 | 560 | 0 | 0 | Luvisol | pasture |
| SEG041_11 | 5.3 | 20.73 | 0 | 20.73 | 1.99 | 10.41 | 136 | 185 | 679 | 1.4 | 362 | 0 | 0 | Luvisol | pasture |
| SEG042_11 | 5.08 | 18.64 | 0 | 18.64 | 1.85 | 10.06 | 117 | 289 | 594 | 2.08 | 801 | 0 | 0 | Luvisol | pasture |
| SEG043_11 | 5.94 | 19.81 | 0.3 | 19.51 | 1.93 | 10.12 | 131 | 121 | 748 | 1.71 | 541 | 0 | 0 | Luvisol | pasture |
| SEG044_11 | 5.41 | 21.83 | 0 | 21.83 | 2.15 | 10.15 | 113 | 189 | 698 | 0.76 | 107 | 0 | 0 | Cambisol | pasture |
| SEG045_11 | 5.26 | 18.91 | 0 | 18.91 | 1.85 | 10.21 | 100 | 234 | 666 | 1.1 | 226 | 0 | 0 | Albeluvisol | pasture |
| SEG046_11 | 5.37 | 24.3 | 0 | 24.3 | 2.3 | 10.58 | 146 | 210 | 644 | 1.77 | 582 | 0 | 0 | Cambisol | pasture |
| SEG047_11 | 6.17 | 23.34 | 0.32 | 23.03 | 2.18 | 10.55 | 186 | 255 | 559 | 2.07 | 599 | 1 | 0 | Luvisol | pasture |
| SEG048_11 | 6.73 | 18.16 | 0.62 | 17.54 | 1.73 | 10.11 | 130 | 120 | 750 | 1.57 | 460 | 0 | 0 | Luvisol | pasture |
| SEG049_11 | 5.98 | 20.98 | 0 | 20.98 | 1.97 | 10.66 | 106 | 372 | 522 | 1.51 | 422 | 0 | 0 | Albeluvisol | pasture |
| SEG050_11 | 5.07 | 23.05 | 0 | 23.05 | 2.13 | 10.84 | 92 | 112 | 796 | 1 | 187 | 0 | 0 | Cambisol | pasture |
| AEG001_17 | 6.78 | 90.96 | 1.41 | 89.56 | 9.19 | 9.75 | 643 | 324 | 33 | 2.32 | 0 | 2 | 30 | Leptosol | meadow |
| AEG002_17 | 6.86 | 82.5 | 2.22 | 80.27 | 8.42 | 9.53 | 512 | 411 | 77 | 13.54 | 0 | 3 | 433 | Leptosol | meadow |
| AEG003_17 | 6.1 | 63.93 | 0 | 63.93 | 5.98 | 10.69 | 672 | 298 | 30 | 2.18 | 23.7 | 2 | 12.92 | Leptosol | meadow |
| AEG004_17 | 5.27 | 51.1 | 0 | 51.1 | 5.26 | 9.71 | 488 | 427 | 85 | 2.52 | 75 | 1 | 30 | Leptosol | mowed_meadow |
| AEG005_17 | 6.26 | 95.35 | 0.68 | 94.67 | 9.39 | 10.08 | 365 | 518 | 117 | 1.55 | 35.9 | 0 | 41.65 | Leptosol | mowed_meadow |
| AEG006_17 | 6.02 | 84.56 | 0.4 | 84.17 | 8.38 | 10.05 | 588 | 380 | 32 | 4.99 | 245 | 1 | 40 | Leptosol | mowed_meadow |
| AEG007_17 | 7.34 | 92.51 | 33.91 | 58.61 | 4.98 | 11.78 | 385 | 427 | 188 | 0.74 | 57.2 | 0 | 0 | Leptosol | pasture |
| AEG008_17 | 6.61 | 91.86 | 0.46 | 91.4 | 8.39 | 10.89 | 673 | 319 | 8 | 1.39 | 47.7 | 1 | 0 | Leptosol | pasture |
| AEG009_17 | 6.58 | 78.6 | 0.53 | 78.07 | 6.46 | 12.08 | 556 | 417 | 27 | 1.57 | 121 | 0 | 0 | Leptosol | pasture |
| AEG010_17 | 5.85 | 94.09 | 0.44 | 93.65 | 9.14 | 10.24 | 647 | 327 | 26 | 1.02 | 19.4 | 1 | 0 | Leptosol | meadow |
| AEG011_17 | 5.36 | 63.06 | 0 | 63.06 | 5.91 | 10.68 | 643 | 327 | 30 | 2.58 | 0 | 2 | 40 | Leptosol | meadow |
| AEG012_17 | 6.63 | 74.58 | 1.52 | 73.06 | 7.11 | 10.28 | 436 | 512 | 52 | 2.32 | 0 | 2 | 30 | Leptosol | meadow |
| AEG013_17 | 6.26 | 76.38 | 0.57 | 75.81 | 7.14 | 10.61 | 643 | 311 | 46 | 3.06 | 0 | 2 | 58.5 | Leptosol | meadow |
| AEG014_17 | 6.6 | 64.81 | 0.65 | 64.16 | 6.36 | 10.09 | 521 | 453 | 26 | 3.09 | 0 | 2 | 60 | Leptosol | meadow |
| AEG015_17 | 5.66 | 62.61 | 0 | 62.61 | 6.27 | 9.98 | 642 | 340 | 18 | 11.7 | 0 | 3 | 362 | Leptosol | meadow |
| AEG016_17 | 5.97 | 72.88 | 0.36 | 72.52 | 7.08 | 10.24 | 528 | 449 | 23 | 2.83 | 159 | 1 | 0 | Leptosol | mowed_meadow |
| AEG017_17 | 6.89 | 65.39 | 1.82 | 63.56 | 6.27 | 10.14 | 618 | 365 | 17 | 1.88 | 0 | 2 | 13.28 | Leptosol | meadow |
| AEG018_17 | 6.94 | 72.9 | 3.75 | 69.15 | 7.01 | 9.87 | 649 | 332 | 19 | 6.24 | 0 | 3 | 151.6 | Leptosol | meadow |
| AEG019_17 | 5.76 | 79.03 | 0.41 | 78.62 | 7.94 | 9.91 | 637 | 327 | 36 | 7 | 480 | 1 | 0 | Leptosol | mowed_meadow |
| AEG020_17 | 6.69 | 89.97 | 0.84 | 89.13 | 9.42 | 9.47 | 597 | 334 | 69 | 2.23 | 172 | 0 | 0 | Leptosol | pasture |
| AEG021_17 | 5.82 | 70.28 | 0.33 | 69.95 | 6.97 | 10.04 | 587 | 391 | 22 | 7.23 | 445 | 1 | 26.4 | Leptosol | pasture |
| AEG022_17 | 5.68 | 56.61 | 0.35 | 56.27 | 5.05 | 11.14 | 708 | 270 | 22 | 1.09 | 24.8 | 1 | 0 | Leptosol | meadow |
| AEG023_17 | 7.05 | 84.93 | 2.82 | 82.11 | 8.44 | 9.73 | 566 | 387 | 47 | 1.87 | 0 | 2 | 12.92 | Leptosol | meadow |
| AEG024_17 | 6.08 | 80.65 | 0.51 | 80.14 | 7.94 | 10.1 | 638 | 329 | 33 | 5.42 | 33 | 3 | 103.5 | Leptosol | mowed_meadow |
| AEG025_17 | 7.19 | 99.43 | 16.9 | 82.52 | 7.37 | 11.2 | 463 | 406 | 131 | 0.93 | 71.3 | 0 | 0 | Leptosol | pasture |
| AEG026_17 | 6.84 | 74.78 | 1.01 | 73.77 | 6.12 | 12.06 | 679 | 241 | 80 | 3.46 | 267 | 0 | 0 | Leptosol | pasture |
| AEG027_17 | 5.97 | 100.86 | 0.47 | 100.39 | 8.72 | 11.51 | 576 | 410 | 14 | 1.42 | 110 | 0 | 0 | Leptosol | pasture |

| | | | | | | | | | | | | | | | |
|---|---|---|---|---|---|---|---|---|---|---|---|---|---|---|---|
| AEG028_17 | 6.13 | 60.01 | 0.33 | 59.69 | 5.21 | 11.46 | 599 | 385 | 16 | 1.11 | 85.9 | 0 | 0 | Leptosol | pasture |
| AEG029_17 | 5.87 | 58.51 | 0.44 | 58.07 | 5.58 | 10.4 | 566 | 343 | 91 | 1.53 | 58.7 | 1 | 0 | Cambisol | mowed_meadow |
| AEG030_17 | 6.64 | 64.4 | 0.56 | 63.85 | 6.47 | 9.86 | 556 | 375 | 69 | 1.18 | 32 | 1 | 0 | Leptosol | mowed_meadow |
| AEG031_17 | 6.66 | 73.28 | 0.68 | 72.6 | 7.25 | 10.01 | 595 | 381 | 24 | 4.59 | 294 | 1 | 0 | Leptosol | mowed_meadow |
| AEG032_17 | 5.41 | 46.34 | 0.3 | 46.04 | 4.12 | 11.16 | 461 | 466 | 73 | 0.78 | 59.8 | 0 | 0 | Leptosol | pasture |
| AEG033_17 | 6 | 50.42 | 0.35 | 50.08 | 4.65 | 10.76 | 597 | 388 | 15 | 0.68 | 52.1 | 0 | 0 | Leptosol | pasture |
| AEG034_17 | 6.33 | 47.3 | 0.43 | 46.87 | 4.72 | 9.93 | 423 | 554 | 23 | 1.66 | 68.8 | 1 | 0 | Leptosol | pasture |
| AEG035_17 | 5.34 | 45.41 | 0.46 | 44.96 | 4.47 | 10.06 | 446 | 503 | 51 | 2.58 | 0 | 2 | 40 | Cambisol | meadow |
| AEG036_17 | 5.97 | 59.44 | 0.5 | 58.94 | 5.58 | 10.57 | 588 | 213 | 199 | 3.72 | 0 | 2 | 84 | Cambisol | meadow |
| AEG037_17 | 6.31 | 73.35 | 0.81 | 72.54 | 6.92 | 10.49 | 566 | 334 | 100 | 2.49 | 0 | 2 | 36.6 | Cambisol | meadow |
| AEG038_17 | 5.62 | 53.76 | 0.29 | 53.47 | 5.51 | 9.7 | 451 | 468 | 81 | 1.96 | 0 | 2 | 16.15 | Cambisol | meadow |
| AEG039_17 | 5.97 | 73.01 | 0.49 | 72.52 | 6.89 | 10.53 | 463 | 493 | 44 | 2.32 | 0 | 2 | 30 | Cambisol | meadow |
| AEG040_17 | 6.86 | 73.61 | 1.7 | 71.91 | 7.36 | 9.76 | 698 | 269 | 33 | 4.66 | 0 | 3 | 90.73 | Cambisol | meadow |
| AEG041_17 | 6.26 | 52.36 | 0.31 | 52.05 | 4.97 | 10.48 | 541 | 407 | 52 | 5.35 | 0 | 2 | 146.9 | Cambisol | meadow |
| AEG042_17 | 7.12 | 99.44 | 16.14 | 83.3 | 8.84 | 9.42 | 159 | 757 | 84 | 3.09 | 0 | 3 | 30 | Cambisol | mowed_meadow |
| AEG043_17 | 6.86 | 90.09 | 4.81 | 85.28 | 8.87 | 9.61 | 613 | 349 | 38 | 1.7 | 71.7 | 1 | 0 | Cambisol | mowed_meadow |
| AEG044_17 | 7.27 | 85.85 | 14.18 | 71.67 | 7.49 | 9.56 | 152 | 735 | 113 | 2.97 | 229 | 0 | 0 | Cambisol | pasture |
| AEG045_17 | 5.39 | 59.82 | 0.32 | 59.5 | 6.1 | 9.75 | 529 | 417 | 54 | 2.41 | 0 | 3 | 3.96 | Cambisol | meadow |
| AEG046_17 | 6.04 | 80.61 | 0.49 | 80.11 | 8.21 | 9.76 | 659 | 287 | 54 | 3.58 | 276 | 0 | 0 | Cambisol | pasture |
| AEG047_17 | 7.51 | 106.49 | 44.73 | 61.75 | 5.65 | 10.92 | 146 | 704 | 150 | 1.25 | 96.4 | 0 | 0 | Cambisol | pasture |
| AEG048_17 | 7.56 | 97.63 | 55.36 | 42.27 | 3.47 | 12.18 | 295 | 565 | 140 | 0.61 | 47.3 | 0 | 0 | Cambisol | pasture |
| AEG049_17 | 6.03 | 56.03 | 0.43 | 55.6 | 5.25 | 10.59 | 451 | 522 | 27 | 1.21 | 93.2 | 0 | 0 | Cambisol | pasture |
| AEG050_17 | 6.04 | 75.07 | 0.48 | 74.59 | 7.41 | 10.06 | 589 | 382 | 29 | 2.39 | 0 | 3 | 3.3 | Cambisol | meadow |
| HEG001_17 | 6.62 | 53.65 | 0.65 | 53 | 5.25 | 10.09 | 502 | 447 | 50 | 10.12 | 112 | 2 | 215.89 | Cambisol | meadow |
| HEG002_17 | 7.26 | 42.16 | 4.97 | 37.19 | 3.88 | 9.59 | 552 | 396 | 52 | 5.95 | 0 | 2 | 130 | Vertisol | meadow |
| HEG003_17 | 7.32 | 45.98 | 7.26 | 38.72 | 4.01 | 9.65 | 544 | 399 | 57 | 6.83 | 0 | 3 | 130 | Vertisol | meadow |
| HEG004_17 | 6.54 | 66.24 | 0.55 | 65.69 | 6.16 | 10.67 | 511 | 414 | 75 | 5.11 | 0 | 2 | 104 | Stagnosol | mowed_meadow |
| HEG005_17 | 7.16 | 46.84 | 4.86 | 41.98 | 4.33 | 9.69 | 454 | 497 | 49 | 5.25 | 0 | 3 | 81 | Stagnosol | mowed_meadow |
| HEG006_17 | 5.85 | 24.42 | 0.28 | 24.14 | 2.39 | 10.11 | 257 | 698 | 45 | 5.33 | 14.4 | 2 | 105.11 | Stagnosol | mowed_meadow |
| HEG007_17 | 7 | 57.5 | 4.29 | 53.21 | 5.6 | 9.5 | 536 | 434 | 30 | 2.58 | 208 | 0 | 0 | Stagnosol | pasture |
| HEG008_17 | 7.01 | 70.05 | 3.06 | 67 | 6.55 | 10.22 | 494 | 452 | 56 | 2.58 | 208 | 0 | 0 | Stagnosol | pasture |
| HEG009_17 | 7.1 | 45.93 | 2.23 | 43.7 | 4.15 | 10.53 | 387 | 537 | 76 | 1.62 | 59.8 | 1 | 0 | Stagnosol | pasture |
| HEG010_17 | 6.54 | 51.95 | 0.44 | 51.51 | 4.56 | 11.29 | 436 | 532 | 30 | 1.65 | 24.9 | 1 | 14.25 | Vertisol | meadow |
| HEG011_17 | 7.28 | 66.85 | 9.92 | 56.92 | 5.41 | 10.52 | 552 | 405 | 43 | 2.11 | 62.3 | 1 | 14.25 | Stagnosol | meadow |
| HEG012_17 | 7.02 | 75.97 | 3.14 | 72.83 | 7.39 | 9.86 | 497 | 448 | 55 | 8.9 | 517 | 1 | 50 | Stagnosol | mowed_meadow |
| HEG013_17 | 7.21 | 48.05 | 4.07 | 43.98 | 4.54 | 9.68 | 394 | 536 | 70 | 5.27 | 48 | 3 | 63.3 | Stagnosol | mowed_meadow |
| HEG014_17 | 6.41 | 44.49 | 0.39 | 44.1 | 4.1 | 10.75 | 450 | 491 | 60 | 2.81 | 0 | 1 | 60 | Stagnosol | mowed_meadow |
| HEG015_17 | 7.07 | 64.42 | 1.77 | 62.66 | 6.03 | 10.38 | 528 | 435 | 38 | 4.34 | 60.1 | 1 | 84 | Stagnosol | mowed_meadow |
| HEG016_17 | 6.76 | 73.31 | 0.83 | 72.48 | 7.07 | 10.24 | 555 | 405 | 39 | 1.39 | 112 | 0 | 0 | Stagnosol | pasture |
| HEG017_17 | 6.93 | 57.03 | 0.62 | 56.41 | 5.18 | 10.9 | 546 | 419 | 32 | 0.77 | 61.7 | 0 | 0 | Stagnosol | pasture |
| HEG018_17 | 7.35 | 65.16 | 11.07 | 54.09 | 4.82 | 11.23 | 457 | 446 | 97 | 1.19 | 25 | 1 | 0 | Vertisol | pasture |
| HEG019_17 | 6.6 | 64.67 | 0.47 | 64.2 | 5.81 | 11.05 | 485 | 427 | 87 | 0.77 | 61.7 | 0 | 0 | Stagnosol | pasture |
| HEG020_17 | 5.52 | 29.73 | 0 | 29.73 | 2.64 | 11.27 | 239 | 661 | 102 | 1.62 | 59.8 | 1 | 0 | Stagnosol | pasture |
| HEG021_17 | 7.3 | 46.68 | 9.42 | 37.25 | 3.6 | 10.35 | 311 | 631 | 58 | 2.01 | 91.6 | 1 | 0 | Stagnosol | pasture |
| HEG022_17 | 6.88 | 53.2 | 1.81 | 51.39 | 5.23 | 9.83 | 446 | 467 | 87 | 2.97 | 97.5 | 2 | 0 | Cambisol | mowed_meadow |
| HEG023_17 | 7.28 | 51.05 | 2.4 | 48.64 | 4.85 | 10.03 | 588 | 375 | 37 | 0.88 | 0 | 1 | 0 | Stagnosol | mowed_meadow |
| HEG024_17 | 6.77 | 61.4 | 1.08 | 60.32 | 6.08 | 9.92 | 545 | 404 | 52 | 1.55 | 54.5 | 1 | 0 | Stagnosol | mowed_meadow |
| HEG025_17 | 7.28 | 62.34 | 3.84 | 58.5 | 5.75 | 10.18 | 423 | 531 | 46 | 1.38 | 40.7 | 1 | 0 | Cambisol | pasture |
| HEG026_17 | 7.37 | 57.04 | 9.92 | 47.12 | 4.85 | 9.72 | 481 | 458 | 61 | 0.88 | 0 | 1 | 0 | Cambisol | meadow |
| HEG027_17 | 7.24 | 55.65 | 4.18 | 51.48 | 4.93 | 10.44 | 492 | 472 | 36 | 3.48 | 53.9 | 1 | 60 | Cambisol | meadow |
| HEG028_17 | 7.32 | 45.17 | 3.63 | 41.54 | 4.21 | 9.87 | 536 | 409 | 55 | 1.72 | 0 | 1 | 26 | Cambisol | mowed_meadow |
| HEG029_17 | 7.16 | 38.63 | 1.01 | 37.62 | 3.6 | 10.46 | 469 | 501 | 30 | 3 | 32.8 | 2 | 26 | Cambisol | mowed_meadow |
| HEG030_17 | 7.19 | 49.27 | 3.07 | 46.21 | 4.77 | 9.68 | 400 | 548 | 52 | 5.69 | 61.9 | 3 | 71 | Cambisol | mowed_meadow |
| HEG031_17 | 7.21 | 53.02 | 6.61 | 46.41 | 4.71 | 9.86 | 237 | 726 | 36 | 2.99 | 50.3 | 1 | 46 | Cambisol | mowed_meadow |
| HEG032_17 | 5.61 | 41.82 | 0.29 | 41.53 | 3.93 | 10.57 | 340 | 640 | 17 | 3.78 | 77.9 | 1 | 60 | Cambisol | mowed_meadow |
| HEG033_17 | 5.34 | 42.08 | 0 | 42.08 | 3.87 | 10.88 | 353 | 618 | 29 | 2.84 | 87.5 | 2 | 0 | Cambisol | pasture |
| HEG034_17 | 6.95 | 40.85 | 1.26 | 39.59 | 4.01 | 9.87 | 448 | 469 | 83 | 5.33 | 14.4 | 2 | 105.11 | Cambisol | mowed_meadow |
| HEG035_17 | 7.02 | 52.99 | 5.33 | 47.65 | 5 | 9.54 | 307 | 580 | 113 | 4.17 | 265 | 1 | 0 | Cambisol | mowed_meadow |
| HEG036_17 | 7.27 | 71.26 | 12.17 | 59.09 | 5.93 | 9.97 | 61 | 871 | 71 | 1.56 | 115 | 0 | 4.28 | Cambisol | mowed_meadow |
| HEG037_17 | 7.3 | 75.89 | 16.62 | 59.27 | 6.18 | 9.58 | 60 | 841 | 99 | 5.04 | 46.1 | 2 | 84 | Cambisol | mowed_meadow |
| HEG038_17 | 7.32 | 36.49 | 5.35 | 31.15 | 3.28 | 9.49 | 310 | 622 | 68 | 4.17 | 336 | 0 | 0 | Cambisol | pasture |
| HEG039_17 | 6.48 | 43.38 | 0.48 | 42.9 | 4.24 | 10.12 | 351 | 537 | 110 | 1.51 | 51.1 | 1 | 0 | Cambisol | pasture |
| HEG040_17 | 6.56 | 68.7 | 1.09 | 67.61 | 6.69 | 10.11 | 275 | 648 | 79 | 4.58 | 369 | 0 | 0 | Cambisol | pasture |
| HEG041_17 | 7.18 | 45.02 | 1.29 | 43.73 | 3.85 | 11.35 | 518 | 417 | 65 | 1.62 | 59.8 | 1 | 0 | Cambisol | pasture |

| | | | | | | | | | | | | | | | |
|---|---|---|---|---|---|---|---|---|---|---|---|---|---|---|---|
| HEG042_17 | 7.24 | 48.67 | 3.85 | 44.82 | 4.2 | 10.68 | 435 | 520 | 46 | 0.77 | 61.7 | 0 | 0 | Cambisol | pasture |
| HEG043_17 | 7.14 | 48.89 | 1.23 | 47.66 | 4.27 | 11.16 | 376 | 589 | 38 | 0.57 | 45.6 | 0 | 0 | Cambisol | pasture |
| HEG044_17 | 7.14 | 87.53 | 4.83 | 82.7 | 8.33 | 9.93 | 79 | 853 | 69 | 0.66 | 53.3 | 0 | 0 | Cambisol | pasture |
| HEG045_17 | 6.98 | 47.47 | 0.64 | 46.83 | 4.49 | 10.42 | 492 | 468 | 43 | 1.45 | 46.1 | 1 | 0 | Cambisol | pasture |
| HEG046_17 | 7.42 | 51.11 | 24.05 | 27.06 | 2.96 | 9.13 | 395 | 531 | 74 | 1.72 | 68.1 | 1 | 0 | Cambisol | pasture |
| HEG047_17 | 7.15 | 62.51 | 2.59 | 59.92 | 5.98 | 10.02 | 483 | 477 | 40 | 2.13 | 101 | 1 | 0 | Cambisol | mowed_meadow |
| HEG048_17 | 7.02 | 48.19 | 0.8 | 47.39 | 4.55 | 10.41 | 465 | 488 | 50 | 1.43 | 44.3 | 1 | 0 | Cambisol | mowed_meadow |
| HEG049_17 | 6.67 | 55.09 | 0.81 | 54.27 | 5.33 | 10.18 | 431 | 500 | 71 | 2.21 | 0 | 2 | 14.25 | Cambisol | mowed_meadow |
| HEG050_17 | 6.89 | 50.6 | 1.02 | 49.58 | 4.71 | 10.52 | 646 | 335 | 23 | 1.75 | 70.6 | 1 | 0 | Cambisol | mowed_meadow |
| SEG001_17 | 7.51 | 190.89 | 17.28 | 173.6 | 20.48 | 8.48 | 233 | 466 | 301 | 2.64 | 24.8 | 2 | 0 | Histosol | meadow |
| SEG002_17 | 7.47 | 148.59 | 43.96 | 104.63 | 13.63 | 7.68 | 198 | 527 | 275 | 2.09 | 149 | 1 | 0 | Histosol | meadow |
| SEG003_17 | 7.6 | 115.62 | 47.34 | 68.29 | 9.29 | 7.35 | 89 | 600 | 311 | 1.9 | 116 | 1 | 0 | Histosol | meadow |
| SEG004_17 | 7.5 | 197.84 | 73.39 | 124.45 | 16.32 | 7.63 | 163 | 686 | 151 | 0 | 0 | 0 | 0 | Histosol | mowed_meadow |
| SEG005_17 | 7.59 | 165.05 | 70.18 | 94.87 | 12.84 | 7.39 | 162 | 575 | 263 | 1.25 | 0 | 1 | 0 | Gleysol | mowed_meadow |
| SEG006_17 | 5.49 | 248.12 | 3.44 | 244.68 | 25.41 | 9.63 | 248 | 284 | 468 | 1.33 | 235 | 0 | 0 | Histosol | mowed_meadow |
| SEG007_17 | 7.48 | 147.4 | 59.9 | 87.5 | 11.58 | 7.55 | 157 | 648 | 195 | 2.52 | 225 | 1 | 0 | Histosol | pasture |
| SEG008_17 | 7.47 | 138.22 | 79.56 | 58.65 | 8.14 | 7.21 | 110 | 713 | 177 | 4.3 | 319 | 2 | 0 | Gleysol | pasture |
| SEG009_17 | 6.63 | 174.98 | 3.92 | 171.06 | 17.1 | 10 | 171 | 143 | 686 | 2.65 | 469 | 0 | 0 | Histosol | pasture |
| SEG010_17 | 7.51 | 220.87 | 59.4 | 161.47 | 17.69 | 9.13 | 183 | 516 | 301 | 1.85 | 105 | 1 | 0 | Histosol | meadow |
| SEG011_17 | 7.54 | 191.72 | 73.26 | 118.45 | 14.87 | 7.97 | 137 | 552 | 311 | 1.85 | 105 | 1 | 0 | Gleysol | meadow |
| SEG012_17 | 7.45 | 135.33 | 40.2 | 95.13 | 12.57 | 7.57 | 138 | 548 | 314 | 2.64 | 24.8 | 2 | 0 | Histosol | meadow |
| SEG013_17 | 5.54 | 21.76 | 0 | 21.76 | 2.4 | 9.08 | 41 | 324 | 635 | 25.71 | 0 | 2 | 10.26 | Cambisol | meadow |
| SEG014_17 | 7.51 | 162.15 | 71.18 | 90.97 | 12.11 | 7.51 | 169 | 625 | 206 | 2.55 | 9.51 | 2 | 0 | Gleysol | pasture |
| SEG015_17 | 7.46 | 167.96 | 59.69 | 108.26 | 14.6 | 7.41 | 191 | 593 | 216 | 1.3 | 8.9 | 1 | 0 | Histosol | mowed_meadow |
| SEG016_17 | 7.43 | 179.48 | 63.07 | 116.42 | 15.34 | 7.59 | 178 | 655 | 167 | 0.14 | 24.1 | 0 | 0 | Gleysol | mowed_meadow |
| SEG017_17 | 5.41 | 320.86 | 4.28 | 316.59 | 30.58 | 10.35 | 374 | 348 | 278 | 2.62 | 243 | 1 | 0 | Histosol | mowed_meadow |
| SEG018_17 | 4.86 | 10.76 | 0 | 10.76 | 1.11 | 9.72 | 65 | 89 | 846 | 1.25 | 0 | 1 | 0 | Luvisol | meadow |
| SEG019_17 | 7.53 | 92.35 | 25.06 | 67.29 | 8.55 | 7.87 | 87 | 553 | 360 | 1.67 | 74.7 | 1 | 0 | Gleysol | pasture |
| SEG020_17 | 6.38 | 246.8 | 6.22 | 240.58 | 24.13 | 9.97 | 307 | 304 | 389 | 2.27 | 402 | 0 | 0 | Histosol | pasture |
| SEG021_17 | 5.26 | 233.04 | 2.48 | 230.56 | 21.72 | 10.61 | 278 | 310 | 412 | 2.51 | 445 | 0 | 0 | Gleysol | pasture |
| SEG022_17 | 7.48 | 95 | 47 | 48 | 6.82 | 7.04 | 93 | 636 | 271 | 0.36 | 64 | 0 | 0 | Gleysol | pasture |
| SEG023_17 | 5.16 | 288.03 | 3.61 | 284.42 | 25.05 | 11.35 | 373 | 461 | 166 | 2.5 | 0 | 2 | 0 | Histosol | meadow |
| SEG024_17 | 7.49 | 221.31 | 35.94 | 185.37 | 23.21 | 7.99 | 283 | 504 | 213 | 1.32 | 11.7 | 1 | 0 | Histosol | meadow |
| SEG025_17 | 6.54 | 183.95 | 4.29 | 179.66 | 17.83 | 10.07 | 176 | 206 | 618 | 2.5 | 0 | 2 | 0 | Histosol | meadow |
| SEG026_17 | 7.23 | 277.07 | 8.45 | 268.62 | 27.08 | 9.92 | 427 | 353 | 220 | 2.5 | 0 | 2 | 0 | Histosol | meadow |
| SEG027_17 | 6.13 | 372.51 | 12.45 | 360.06 | 30.64 | 11.75 | 416 | 417 | 167 | 1.25 | 0 | 1 | 0 | Histosol | meadow |
| SEG028_17 | 7.52 | 189.8 | 57.87 | 131.93 | 16.09 | 8.2 | 175 | 651 | 174 | 1.31 | 11.2 | 1 | 0 | Histosol | meadow |
| SEG029_17 | 7.61 | 163.06 | 53.21 | 109.85 | 14.39 | 7.63 | 195 | 694 | 110 | 0.5 | 87.7 | 0 | 0 | Histosol | meadow |
| SEG030_17 | 7.04 | 29.35 | 1.77 | 27.59 | 3.17 | 8.71 | 146 | 343 | 511 | 1.25 | 0 | 1 | 0 | Albeluvisol | meadow |
| SEG031_17 | 5.95 | 27.5 | 0 | 27.5 | 2.69 | 10.21 | 182 | 281 | 537 | 1.25 | 0 | 1 | 0 | Cambisol | meadow |
| SEG032_17 | 5.65 | 22.55 | 0 | 22.55 | 2.21 | 10.21 | 190 | 234 | 576 | 1.25 | 0 | 1 | 0 | Luvisol | meadow |
| SEG033_17 | 5.69 | 20.09 | 0 | 20.09 | 1.98 | 10.15 | 118 | 265 | 617 | 1.67 | 295 | 0 | 0 | Albeluvisol | mowed_meadow |
| SEG034_17 | 5.76 | 20.45 | 0 | 20.45 | 1.92 | 10.67 | 138 | 273 | 589 | 2.34 | 192 | 1 | 0 | Albeluvisol | mowed_meadow |
| SEG035_17 | 6.18 | 25.23 | 0 | 25.23 | 2.43 | 10.37 | 200 | 238 | 562 | 0.97 | 171 | 0 | 0 | Luvisol | mowed_meadow |
| SEG036_17 | 6.29 | 24.61 | 0 | 24.61 | 2.36 | 10.42 | 139 | 301 | 560 | 2.1 | 151 | 1 | 0 | Albeluvisol | pasture |
| SEG037_17 | 4.71 | 14.37 | 0 | 14.37 | 1.31 | 10.95 | 93 | 99 | 808 | 0.45 | 79.9 | 0 | 0 | Albeluvisol | pasture |
| SEG038_17 | 5.15 | 18.47 | 0 | 18.47 | 1.71 | 10.8 | 89 | 72 | 838 | 6.48 | 925 | 1 | 0 | Cambisol | mowed_meadow |
| SEG039_17 | 7.4 | 25.96 | 6.22 | 19.74 | 1.95 | 10.13 | 225 | 237 | 538 | 2.89 | 68.8 | 2 | 0 | Cambisol | pasture |
| SEG040_17 | 6.12 | 45.64 | 0 | 45.64 | 4.39 | 10.4 | 98 | 192 | 710 | 1.05 | 186 | 0 | 0 | Luvisol | pasture |
| SEG041_17 | 6.11 | 24.51 | 0 | 24.51 | 2.37 | 10.34 | 136 | 185 | 679 | 3.11 | 330 | 1 | 0 | Luvisol | pasture |
| SEG042_17 | 5.01 | 21.09 | 0 | 21.09 | 2.08 | 10.15 | 117 | 289 | 594 | 2.99 | 528 | 0 | 0 | Luvisol | pasture |
| SEG043_17 | 6.51 | 29.27 | 0.36 | 28.91 | 2.77 | 10.42 | 131 | 121 | 748 | 28.14 | 239 | 0 | 11.84 | Luvisol | pasture |
| SEG044_17 | 5.45 | 21.63 | 0 | 21.63 | 2.1 | 10.3 | 113 | 189 | 698 | 0.96 | 169 | 0 | 0 | Cambisol | pasture |
| SEG045_17 | 5.89 | 18.83 | 0 | 18.83 | 1.79 | 10.51 | 100 | 234 | 666 | 1.01 | 179 | 0 | 0 | Albeluvisol | pasture |
| SEG046_17 | 7.07 | 38.57 | 4.1 | 34.48 | 3.59 | 9.61 | 146 | 210 | 644 | 5.07 | 676 | 1 | 0 | Cambisol | pasture |
| SEG047_17 | 5.7 | 26.15 | 0 | 26.15 | 2.37 | 11.02 | 186 | 255 | 559 | 2.5 | 221 | 1 | 0 | Luvisol | pasture |
| SEG048_17 | 6.67 | 15.81 | 0.34 | 15.47 | 1.56 | 9.89 | 130 | 120 | 750 | 2.95 | 521 | 0 | 0 | Luvisol | pasture |
| SEG049_17 | 6.36 | 22.33 | 0.37 | 21.96 | 2.15 | 10.22 | 106 | 372 | 522 | 2.14 | 378 | 0 | 0 | Albeluvisol | pasture |
| SEG050_17 | 5.44 | 24.83 | 0 | 24.83 | 2.19 | 11.36 | 92 | 112 | 796 | 2.15 | 380 | 0 | 0 | Cambisol | pasture |

1: (cattle*day)/year
2: (cuts/year)
