## Supplementary material for "Contrasting responses of protistan plant parasites and phagotrophs to ecosystems, land management and soil properties": Table S5

**Table S5**. Most parsimonious models (dbRDA), with their respective $R^2$ adjusted and F values, and the F values of the factors selected by each model. Significance values shown as symbol (see footnote).

### CERCOZOA & ENDOMYXA

| Grassland | $R^2$ (%) | Anova F value | soil type | pH | clay | manage-ment | LUI | fertili-zation | mowing | C/N ratio |
|---|---|---|---|---|---|---|---|---|---|---|
| **All OTUs** | 5.5 | 2.4 *** | 2.3 ** | | | 1.6 * | 2.0 ** | 2.0 ** | | 3.5 ** |
| **Bacterivores** | 4.2 | 2.6 *** | 2.6 * | | | | | | | 2.9 * |
| **Omnivores** | 5.2 | 2.6 *** | 2.6 ** | | | 1.7 * | | | | 4.3 ** |
| **Eukaryvores** | 6.0 | 2.8 *** | 2.7 ** | | | 2.5 ** | | | | 4.9 ** |
| **Plant parasites** | 6.3 | 3.2 *** | 3.2 ** | | | 2.5 ** | | | | |

| Grassland by region | $R^2$ (%) | Anova F value | soil type | pH | clay | manage-ment | LUI | fertili-zation | mowing | C/N ratio |
|---|---|---|---|---|---|---|---|---|---|---|
| **Alb** | NS | NS | | | | | | | | |
| **Hainich** | 7.3 | 3.0 *** | 3.6 *** | | | | 2.9 ** | | 4.0 *** | |
| **Schorfheide** | 3.6 | 1.6 ** | 1.6 ** | | 2.1 ** | | 1.9 * | | | 2.6 ** |

| Forest | $R^2$ (%) | Anova F value | soil type | pH | clay | main tree species | intensity manag. | develop-mental stage | organic C | C/N ratio |
|---|---|---|---|---|---|---|---|---|---|---|
| **All OTUs** | 47.7 | 23.1 *** | 3.8 *** | 17.1 *** | 9.3 *** | 5.8 *** | | | | 6.0 ** |
| **Bacterivores** | 44.4 | 18.9 *** | 3.7 ** | 10.8 ** | 7.2 ** | 5.5 ** | | | 2.6 * | 5.5 ** |
| **Omnivores** | 53.2 | 28.7 *** | 3.5 ** | 24.9 ** | 8.9 ** | 6.3 ** | | | | 6.2 ** |
| **Eukaryvores** | 42.6 | 19.0 *** | 4.1 ** | 8.1 ** | 8.2 ** | 4.9 ** | | | | 4.9 ** |

| Forest by region | $R^2$ (%) | Anova F value | soil type | pH | clay | main tree species | intensity manag. | develop-mental stage | organic C | C/N ratio |
|---|---|---|---|---|---|---|---|---|---|---|
| **Alb** | 25.9 | 7.9 *** | | 19.9 *** | 4.0 ** | 4.8 *** | | | 2.4 * | 8.5 *** |
| **Hainich** | 10.2 | 4.4 *** | | 4.6 ** | 4.0 ** | 1.9 * | | | 5.3 ** | |
| **Schorfheide** | 28.7 | 4.2 *** | 1.5 ** | 12.9 ** | 1.8 * | 3.6 ** | | 2.1 ** | 3.2 * | 2.7 ** |

*p values signification codes: *** ≤ 0.001; ** ≤ 0.01; * ≤ 0.05.*
