## Supplementary material for "Contrasting responses of protistan plant parasites and phagotrophs to ecosystems, land management and soil properties": Fig. S2

**Figure S2**. Description of the diversity. **A**. Rarefaction curve describing the observed number of OTUs as a function of the sequencing effort; saturation was reached with c. 235,000 sequences. **B**. Species accumulation curve describing the sampling effort; saturation was reached with 65 samples.

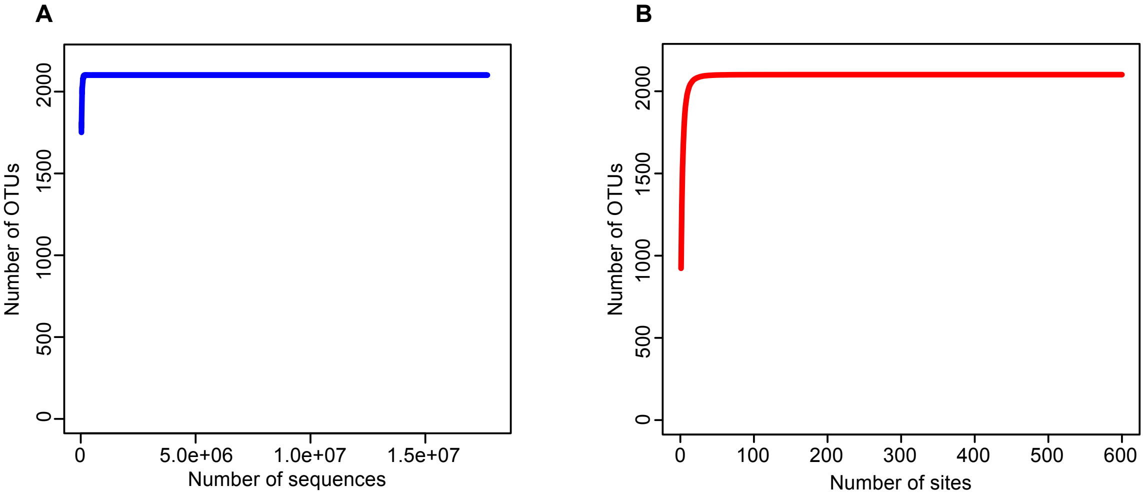
