## Supplementary material for "Contrasting responses of protistan plant parasites and phagotrophs to ecosystems, land management and soil properties": Fig. S3

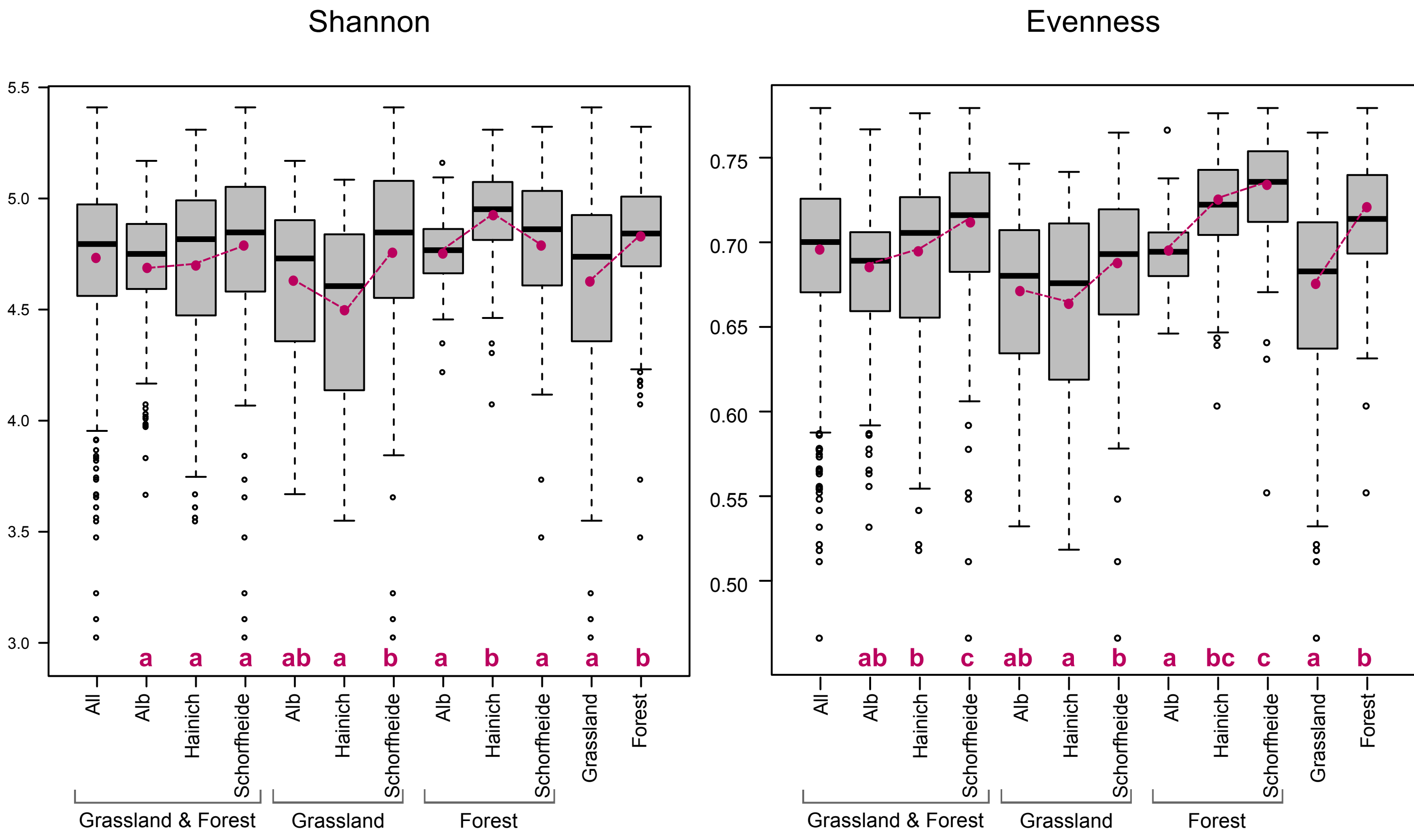


| | All sites | Alb | Hainich | Schorfheide | Alb grassland | Hainich grassland | Schorfheide grassland | Alb forest | Hainich forest | Schorfheide forest | Grassland | Forest |
|---|---|---|---|---|---|---|---|---|---|---|---|---|
| **Shannon** | | | | | | | | | | | | |
| **Min.** | 3.03 | 3.67 | 3.55 | 3.03 | 3.67 | 3.55 | 3.03 | 4.22 | 4.08 | 3.48 | 3.03 | 3.48 |
| **Mean** | 4.72 | 4.70 | 4.70 | **4.77** | 4.63 | 4.49 | **4.76** | 4.76 | **4.92** | 4.79 | 4.63 | **4.82** |
| **Max.** | 5.41 | 5.17 | 5.31 | 5.41 | 5.17 | 5.08 | 5.41 | 5.16 | 5.31 | 5.32 | 5.41 | 5.32 |
| **St_Dev** | 0.36 | 0.27 | 0.40 | 0.40 | 0.34 | 0.41 | 0.46 | 0.17 | 0.22 | 0.33 | 0.42 | 0.26 |
| **evenness** | | | | | | | | | | | | |
| **Min.** | 0.47 | 0.54 | 0.52 | 0.47 | 0.54 | 0.52 | 0.47 | 0.65 | 0.61 | 0.56 | 0.47 | 0.56 |
| **Mean** | 0.70 | 0.68 | 0.69 | **0.71** | 0.67 | 0.66 | **0.69** | 0.70 | 0.72 | **0.73** | 0.67 | **0.72** |
| **Max.** | 0.78 | 0.77 | 0.78 | 0.78 | 0.75 | 0.75 | 0.77 | 0.77 | 0.78 | 0.78 | 0.77 | 0.78 |
| **St_Dev** | 0.05 | 0.04 | 0.06 | 0.05 | 0.05 | 0.06 | 0.05 | 0.02 | 0.03 | 0.04 | 0.05 | 0.03 |

**Figure S3**. Boxplots and table of the alpha diversity of the cercozoan and endomyxan OTUs estimated with the Shannon and evenness indices, for all sites and for sites binned by region and ecotype. Red letters: a change from “a” to “b”, or “c” indicates a significant difference (multiple comparison of means, Tukey's test); two or three letters (e.g. “ab” or "abc") indicate non-significant differences between plots sharing those letters. Red dots indicate the means (black lines the median). In the table, the highest means are in bold.
