## Supplementary material for "Contrasting responses of protistan plant parasites and phagotrophs to ecosystems, land management and soil properties": Fig. S4

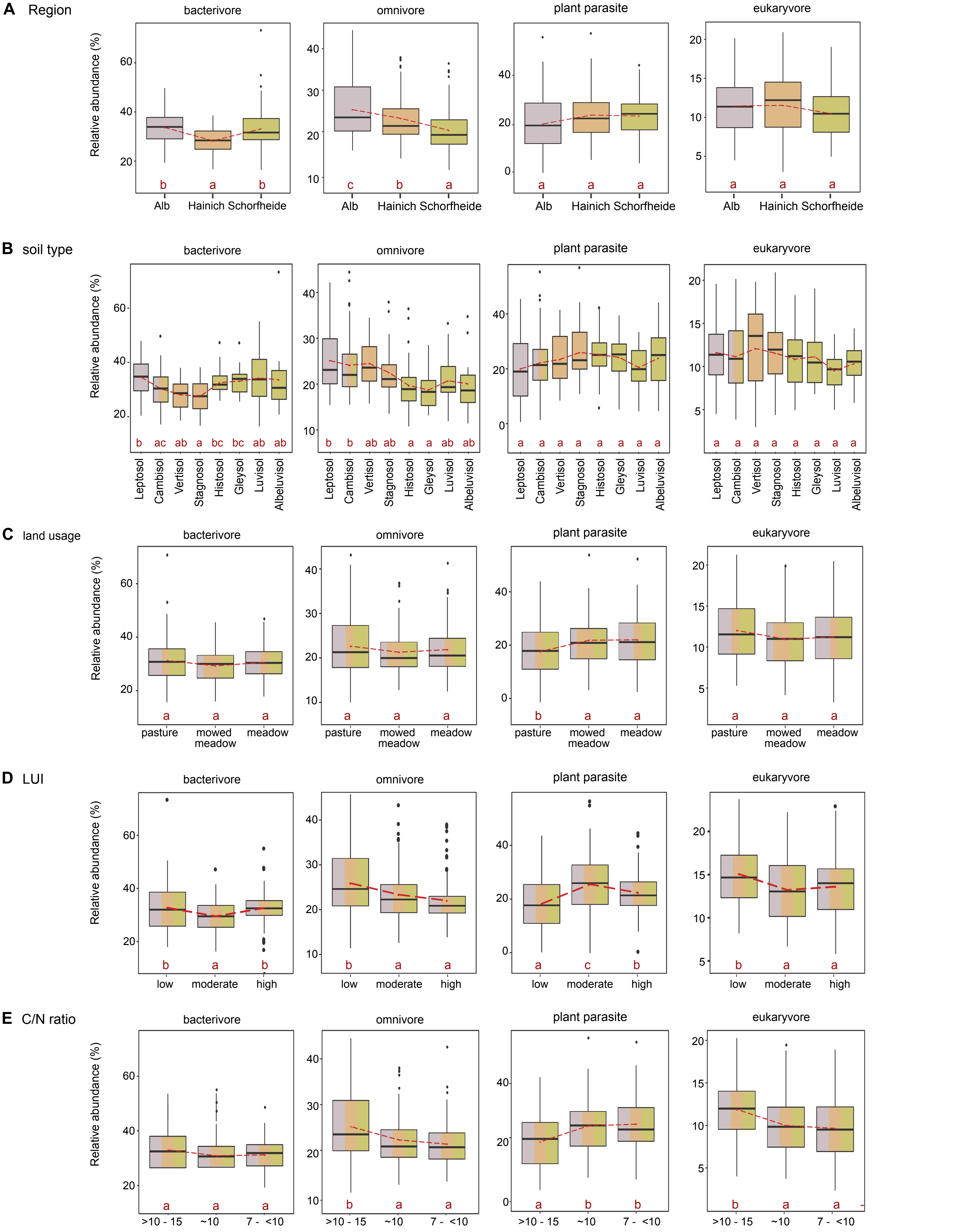


**Figure S4**. Boxplots of the variation of the relative abundances of the four main nutrition modes of Cercozoa and Endomyxa in grassland, colored according to region (approx. %). **A**. by region; **B**. by soil type, only Cambisoil occurs in the three regions; **C**. by grassland management; **D**. by land use intensity (LUI) index, transformed into a categorical variable according to quantiles. **E**. by C/N ratio, transformed as in D. The y-scale varies between graphs. Red letters: a change from "a" to "b", or "c" indicates a significant difference (multiple comparison of means, Tukey's test); two or three letters (e.g. "ab" or "abc") indicate non-significant differences between plots sharing those letters. Red lines indicate the mean. Right side, tables with the regional repartion of each factor; 100 sites in Alb, 96 in Hainich and 97 in Schorfheide.
