## Supplementary material for "Contrasting responses of protistan plant parasites and phagotrophs to ecosystems, land management and soil properties": Fig. S5

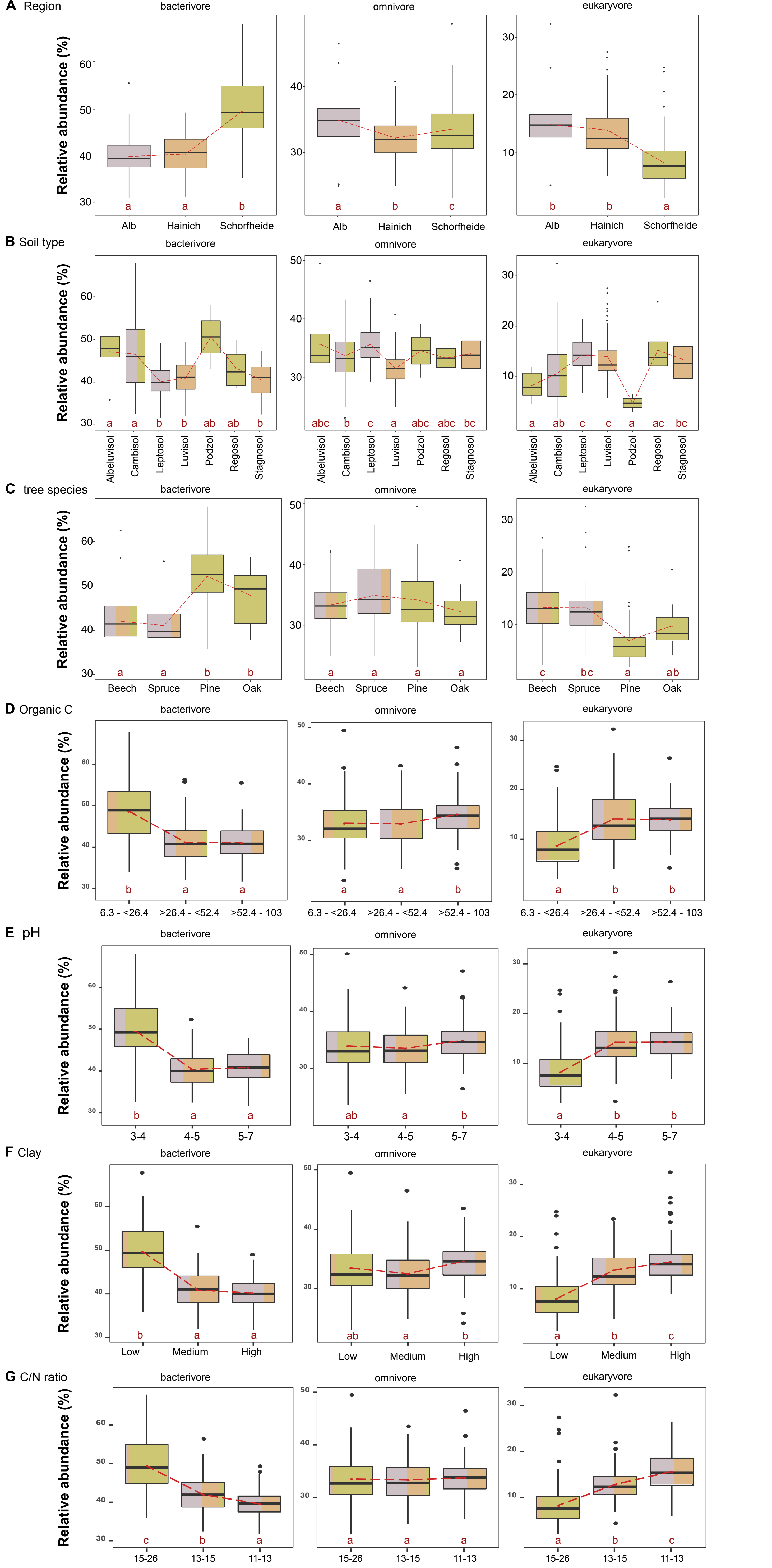


**Figure S5.** Boxplots of the relative abundances of the three main nutrition modes of Cercozoa in forest (nearly no plant parasites were found in forest), coloured according to region. **A**. by region; **B**. by soil type, only Cambisoil is found in two regions; **C**. by main tree species, Latin names: beech = *Fagus sylvatica*, spruce = *Picea abies*, pine = *Pinus sylvestris*, oak = *Quercus petrea* & *Q. robur*, pine and oak growing only in Schorfheide; **D**. by organic carbon (g/kg soil) levels; **E**. by pH levels; **F.** by levels of percentage of soil clay content; **G**. by levels of C/N ratio. The y-scale varies between graphs. Red letters: a change from “a” to “b”, or “c” indicates a significant difference (multiple comparison of means, Tukey's test); two or three letters (e.g. “ab” or "abc") indicate non-significant differences between boxes sharing those letters. Red lines indicate the mean. Right side, tables with regional repartition of each factor; 100 sites in Alb, 95 in Hainich and 97 in Schorfheide.
